# HbtR, a heterofunctional homolog of the virulence regulator TcpP, facilitates the transition between symbiotic and planktonic lifestyles in *Vibrio fischeri*

**DOI:** 10.1101/2020.03.31.019356

**Authors:** Brittany D. Bennett, Tara Essock-Burns, Edward G. Ruby

## Abstract

The bioluminescent bacterium *Vibrio fischeri* forms a mutually beneficial symbiosis with the Hawaiian bobtail squid, *Euprymna scolopes*, in which the bacteria, housed inside a specialized light organ, produce light used by the squid in its nocturnal activities. Upon hatching, *E. scolopes* juveniles acquire *V. fischeri* from the seawater through a complex process that requires, among other factors, chemotaxis by the bacteria along a gradient of *N*-acetylated sugars into the crypts of the light organ, the niche in which the bacteria reside. Once inside the light organ, *V. fischeri* transitions into a symbiotic, sessile state in which the quorum-signaling regulator LitR induces luminescence. In this work we show that expression of *litR* and luminescence are repressed by a homolog of the *V. cholerae* virulence factor TcpP, which we have named HbtR. Further, we demonstrate that LitR represses genes involved in motility and chemotaxis into the light organ and activates genes required for exopolysaccharide production.

**Importance:** TcpP homologs are widespread throughout the *Vibrio* genus; however, the only protein in this family described thus far is a *V. cholerae* virulence regulator. Here we show that HbtR, the TcpP homolog in *V. fischeri*, has both a biological role and regulatory pathway completely unlike that in *V. cholerae*. Through its repression of the quorum-signaling regulator LitR, HbtR affects the expression of genes important for colonization of the *E. scolopes* light organ. While LitR becomes activated within the crypts, and upregulates luminescence and exopolysaccharide genes and downregulates chemotaxis and motility genes, it appears that HbtR, upon expulsion of *V. fischeri* cells into seawater, reverses this process to aid the switch from a symbiotic to a planktonic state. The possible importance of HbtR to the survival of *V. fischeri* outside of its animal host may have broader implications for the ways in which bacteria transition between often vastly different environmental niches.

## Introduction

The Gram-negative bacterium *Vibrio (Aliivibrio) fischeri* is a model organism for the study of biochemical processes underpinning bioluminescence, quorum sensing, and bacterial-animal symbioses. Luminescence in *V. fischeri* is activated by a quorum-sensing pathway that responds to high concentrations of two autoinducer molecules, *N*- (3-oxohexanoyl)-L-homoserine lactone (3-oxo-C6-HSL (1)) and *N*-octanoyl-L-homoserine lactone (C8-HSL (2)). When *V. fischeri* cells multiply to a certain density, 3-oxo-C6-HSL and C8-HSL accumulate to high local concentrations, initiating a signaling cascade that leads to upregulation of the regulator LuxR by the regulator LitR and consequent activation of the luciferase operon, *luxICDABEG* (3–5); reviewed in (6).

Light production by *V. fischeri* is a crucial factor in the mutualistic symbiosis it forms within the light-emitting organ of the Hawaiian bobtail squid, *Euprymna scolopes* (7–9). Juvenile *E. scolopes* become colonized with *V. fischeri* shortly after hatching into seawater containing this bacterium (10). *V. fischeri* cells initiate light-organ colonization through a series of steps, including chemotaxis towards *N*-acetylated sugars released by the squid (11, 12). After initial colonization of the squid light organ, the symbiosis undergoes a daily cyclic rhythm of three basic stages for the remainder of the squid’s life: during the day, *V. fischeri* cells grow to a high density in the crypts on carbon sources provided by the squid (13, 14); at night, the bacteria produce light that aids in camouflage for the squid (15, 16); and at dawn, ∼95% of the bacterial cells are expelled from the light organ into the seawater, where they may initiate colonization of new squid hatchlings, while the remaining ∼5% repopulate the light organ (17, 18).

Previous work indicated that, in *V. fischeri* cells newly expelled from the light organ, numerous genes are up- and downregulated relative to their level of expression in cells that had been planktonic for some time (19). Two of the genes upregulated in expelled cells were VF_A0473 and VF_A0474, comprising a small operon annotated as encoding homologs of the genetic regulator TcpP and its chaperone, TcpH. TcpP was first described in *V. cholerae* as a virulence gene regulator (20, 21). During the early stages of *V. cholerae* infection, TcpP works synergistically with another transcriptional regulator, ToxR, to upregulate expression of the virulence regulator *toxT* and thereby activate expression of cholera toxin and the toxin-coregulated pilus in response to changing environmental conditions (21, 22). *tcpPH* expression is itself induced by AphA and AphB in response to low oxygen or acidic pH (23–25). In *V. cholerae* El Tor biotypes, *aphA* expression is repressed by HapR (the *V. cholerae* ortholog of LitR) upon initiation of quorum sensing at high cell densities (26, 27). Additionally, activation of *tcpPH* expression by AphA and AphB is inhibited by the global metabolic regulator Crp (28). This complex regulatory pathway serves to upregulate *tcpPH* and, consequently, downstream virulence factors under conditions consistent with entry into the vertebrate gut, a process reversed late in the infection cycle before the bacteria exit the host and re-enter an aquatic environment (29, 30).

No published genomes of *V. fischeri* strains encode homologs of either *toxT* or cholera toxin genes. The genome of the model strain *V. fischeri* ES114 does encode annotated homologs of the toxin-coregulated pilus genes *tcpFETSD* and *tcpCQBA*, but this strain is alone among the 66 currently available *V. fischeri* genomes to do so. The *V. fischeri* genes regulated by the protein product of VF_A0473 are therefore unknown, as are any upstream regulatory factors that determine under which conditions the regulator is produced and active. Here we present evidence that VF_A0473 regulates genes governing phenotypes relevant to the switch from symbiosis to a planktonic lifestyle. We therefore rename VF_A0473 and VF_A0474, currently annotated as *tcpP* and *tcpH*, as *hbtR* (*h*a*b*itat *t*ransition *r*egulator) and *hbtC* (*h*a*b*itat *t*ransition *c*haperone). In this work, we identify the HbtR regulon and uncover aspects of *hbtRC* regulation. Further, we determine that LitR is repressed by HbtR and describe LitR-regulated phenotypes beyond luminescence that are important for light-organ colonization.

## Results

### The HbtR regulon is distinct from that of TcpP

Considering that *V. fischeri* and *V. cholerae* have significantly different lifestyles, as well as the absence in 65 of 66 sequenced *V. fischeri* genomes of homologs of virulence-factor genes regulated by TcpP in *V. cholerae*, we postulated that HbtR has a distinct function from that of TcpP. To determine the regulon of HbtR, RNA-seq was performed using Δ*hbtRC* mutant strains of *V. fischeri* ES114 carrying either empty vector or an inducible vector with *hbtRC* under control of the *lac* promoter. Notable among the RNA-seq results (Table S1) is that none of the *tcp* pilus gene homologs encoded in the *V. fischeri* ES114 genome were expressed at significantly different levels in *hbtRC* strains carrying either empty vector or the inducible *hbtRC* vector (Table S1), ruling out the possibility that HbtR regulates the same genes in *V. fischeri* ES114 as TcpP does in *V. cholerae*.

### HbtR represses *litR* expression and luminescence *in vitro*

Among the genes significantly downregulated in the RNA-seq results when *hbtRC* was expressed were the quorum signaling-regulated genes *litR, luxR, qsrP*, and *rpoQ* (Table 1, Table S1). Less strongly, but statistically significantly, downregulated genes included most of the luciferase operon (Table 1, Table S1). To determine whether repression of *lux* genes by HbtR affects bacterial light emission, luminescence and growth were monitored in cultures of wild-type *V. fischeri* and its Δ*hbtRC* derivative carrying either empty vector or vector constitutively expressing *hbtRC*. Deletion of *hbtRC* did not affect light production; however, overexpression of *hbtRC* in either wild type or Δ*hbtRC* led to a delay in onset and significant decrease in intensity of light production (Fig. 1A). There was no difference in growth rate between the strains (Fig. 1B), eliminating the possibility that the reduced luminescence by the *hbtRC*-overexpression strains was due to a growth defect. To determine whether deletion of *hbtRC* would lead to a change in light production by *V. fischeri* within the light organ, *E. scolopes* hatchlings were colonized with wild-type *V. fischeri* or Δ*hbtRC*. There was no difference in the luminescence of juvenile squid colonized by either *V. fischeri* strain at either 24 or 48 h (Fig. S1), suggesting that HbtR is not active inside the light organ.

**Table 1.**
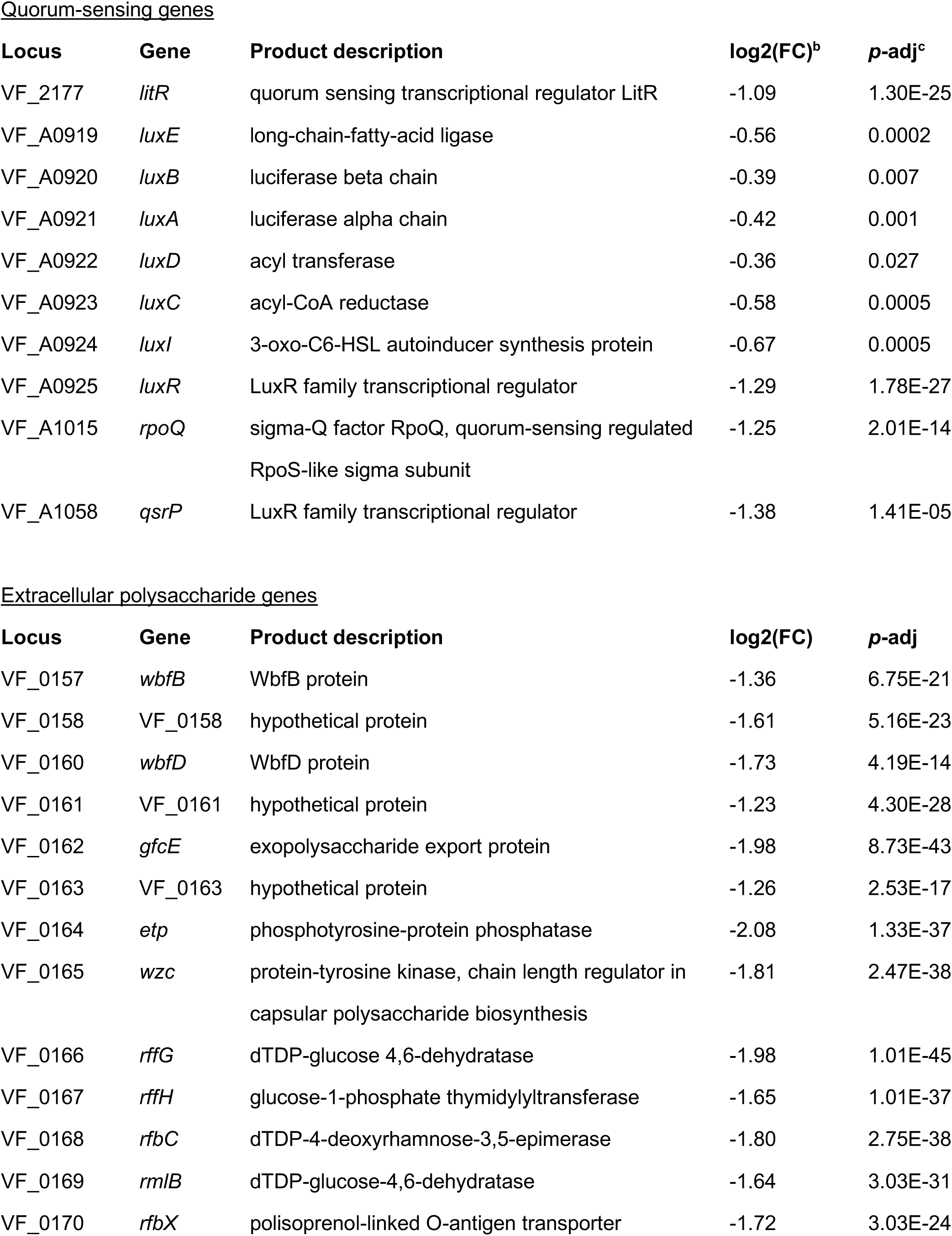
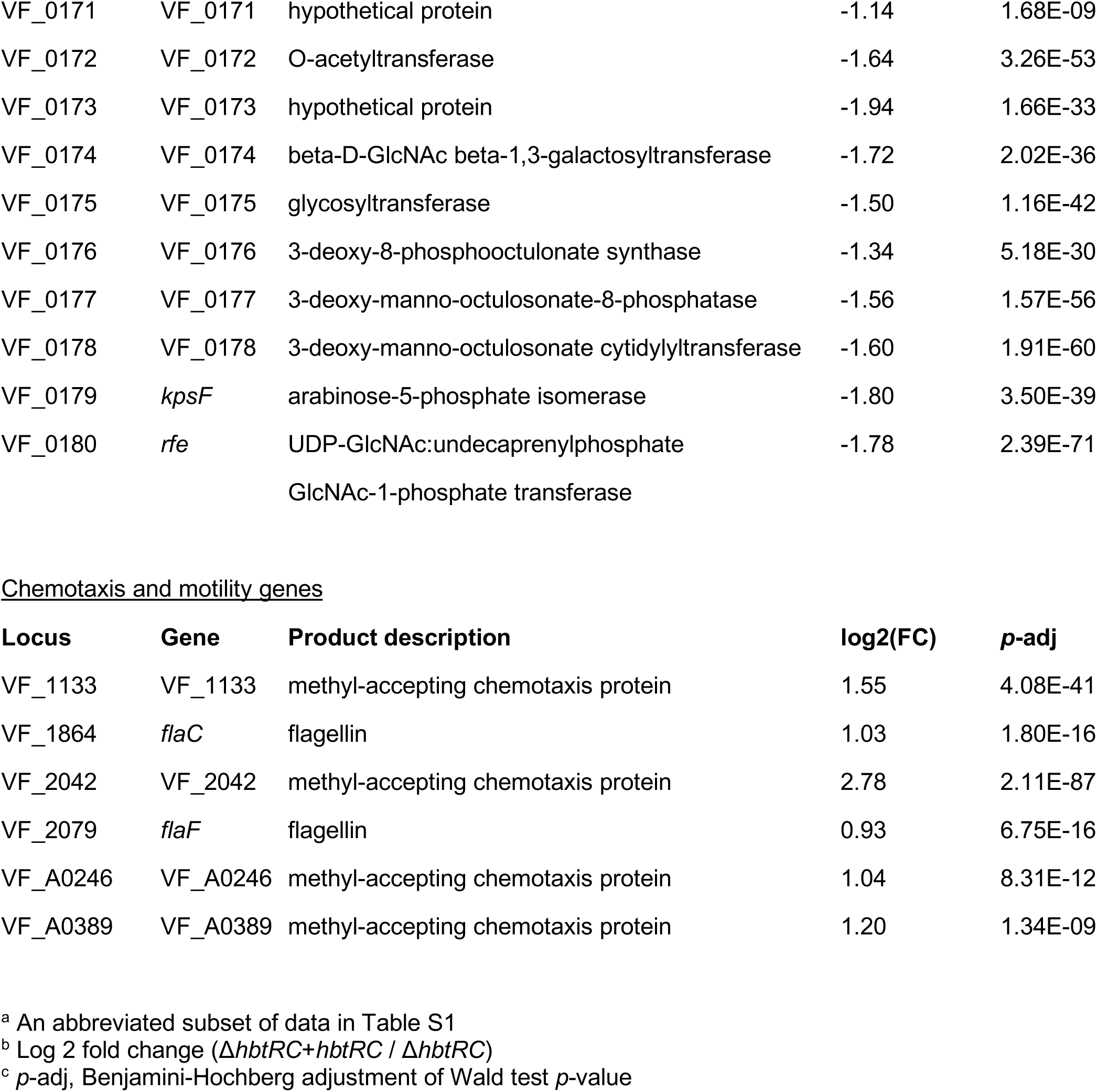
RNA-seq results for genes analyzed in this study^a^.

**FIG 1.**
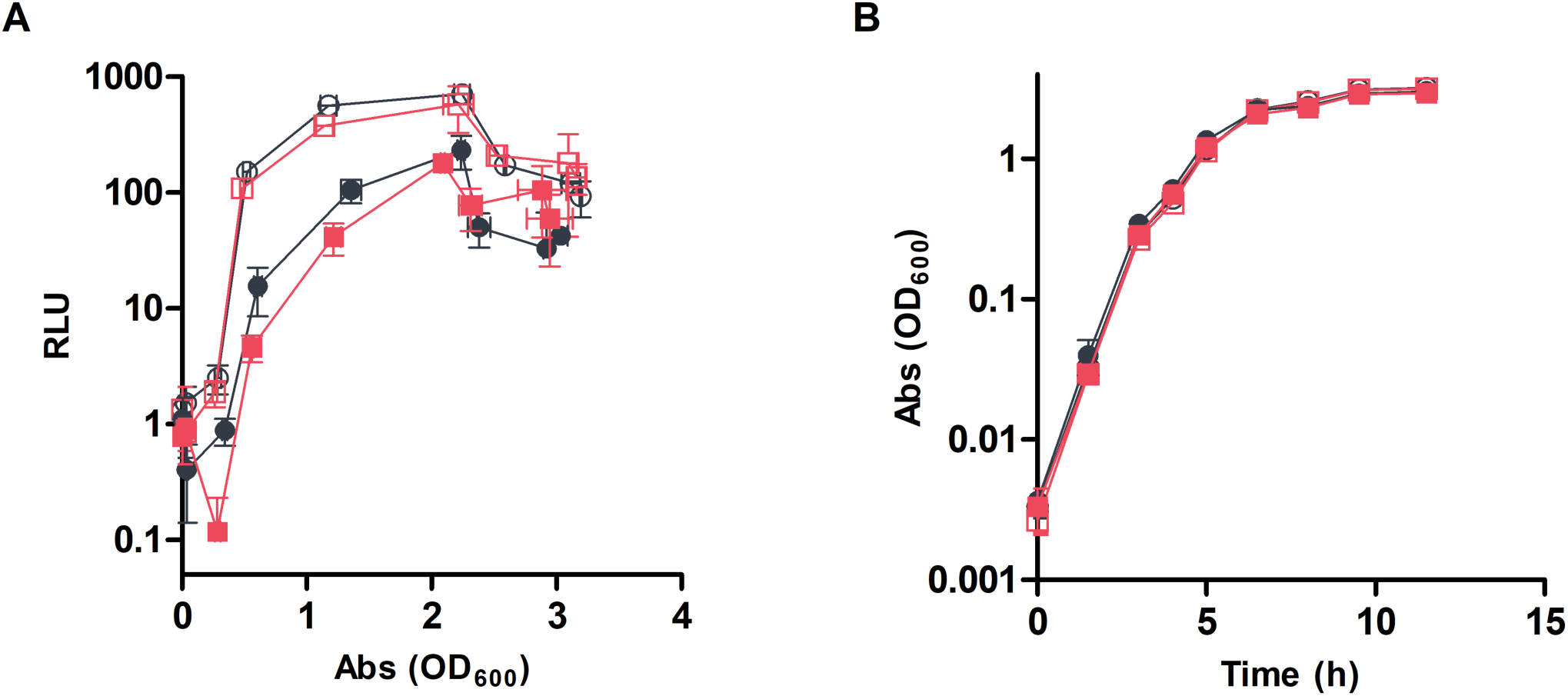
Luminescence and growth of wild-type *V. fischeri* and Δ*hbtRC* strains. The rates of (A) light production and (B) growth in SWT were measured for wild-type *V. fischeri* (circles) and Δ*hbtRC* (squares) carrying empty pVSV105 (open symbols) or pVSV105::*hbtRC* (closed symbols). Results represent the means of three biological replicates ± one standard deviation. RLU, relative light units; Abs, absorbance.

### HbtR activates and LitR represses chemosensory genes involved in *E. scolopes* light-organ colonization

Among the genes upregulated in the *hbtRC*-expressing strain in the RNA-seq experiment were four genes annotated as encoding methyl-accepting chemotaxis proteins (MCPs): VF_1133, VF_2042, VF_A0246, and VF_A0389 (Table 1, Table S1). To confirm that the upregulation of these MCP genes by HbtR functions through its repression of *litR* expression, RT-qPCR was performed on wild-type *V. fischeri* and Δ*litR*, as well as Δ*hbtRC* carrying either empty vector or vector constitutively expressing *hbtRC*. Expression of all four of these MCP genes was significantly increased in Δ*litR* compared to wild-type *V. fischeri* (Table 2). Expression of VF_2042 was significantly increased in Δ*hbtRC* constitutively expressing *hbtRC* compared to Δ*hbtRC* carrying empty vector (Table 2), confirming the RNA-seq results indicating that HbtR affects MCP gene regulation.

**Table 2.**
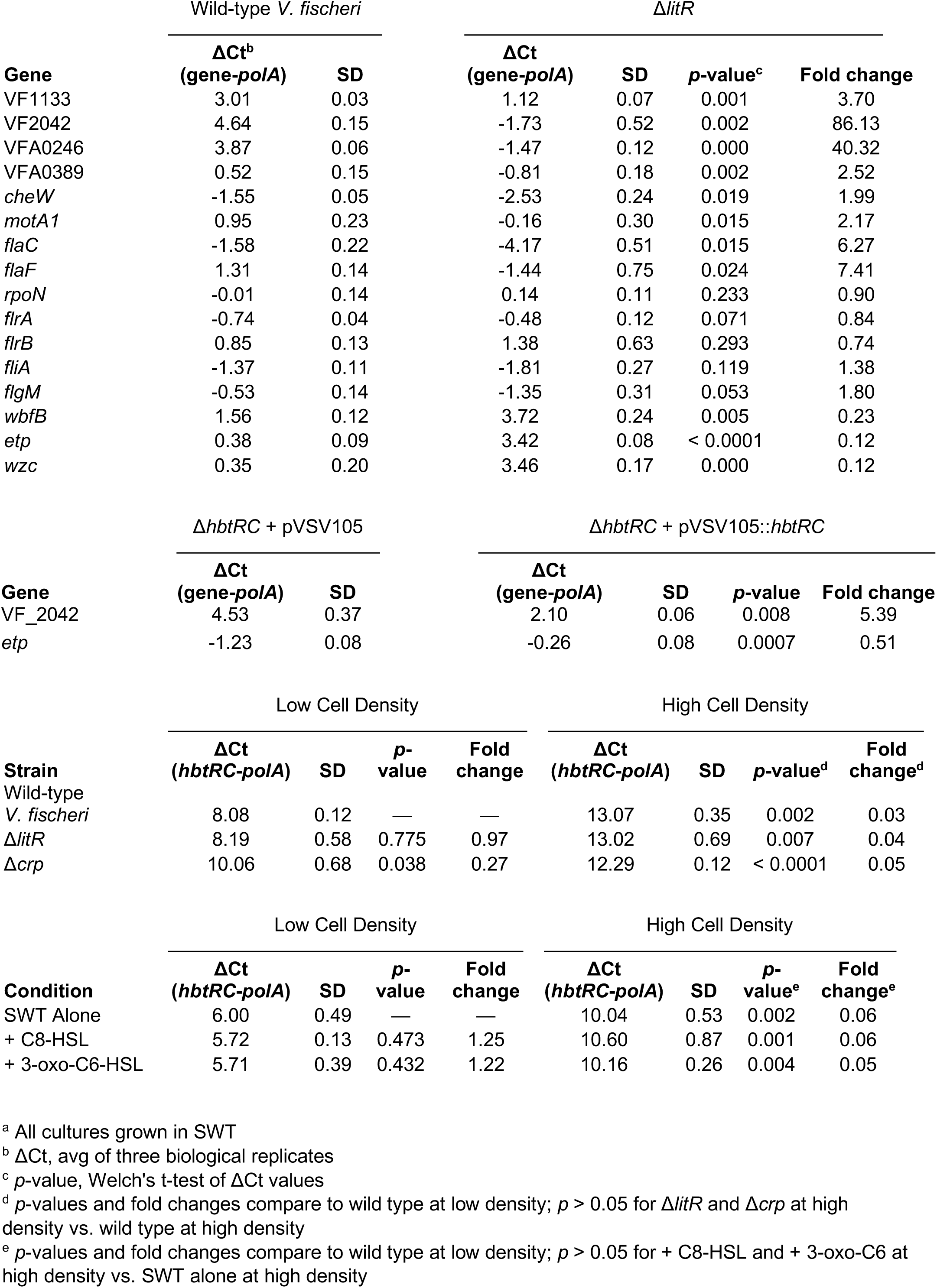
RT-qPCR results^a^.

As chemotaxis is essential to colonization of the *E. scolopes* light organ by *V. fischeri* (12), we asked whether any of the four MCPs repressed by LitR are involved in colonization. Newly hatched *E. scolopes* juveniles were exposed to a 1:1 ratio of wild-type *V. fischeri* and a quadruple mutant with in-frame deletions of all four of the LitR-repressed MCP genes (ΔVF_1133ΔVF_2042ΔVF_A0246ΔVF_A0389; “ΔΔΔΔ”). Regardless of which strain carried a *gfp*-labeled plasmid, the squid light organ populations tended to be dominated by wild-type *V. fischeri* (Fig. 2), indicating that HbtR and LitR regulate chemotaxis genes involved in initiating symbiosis.

**FIG 2.**
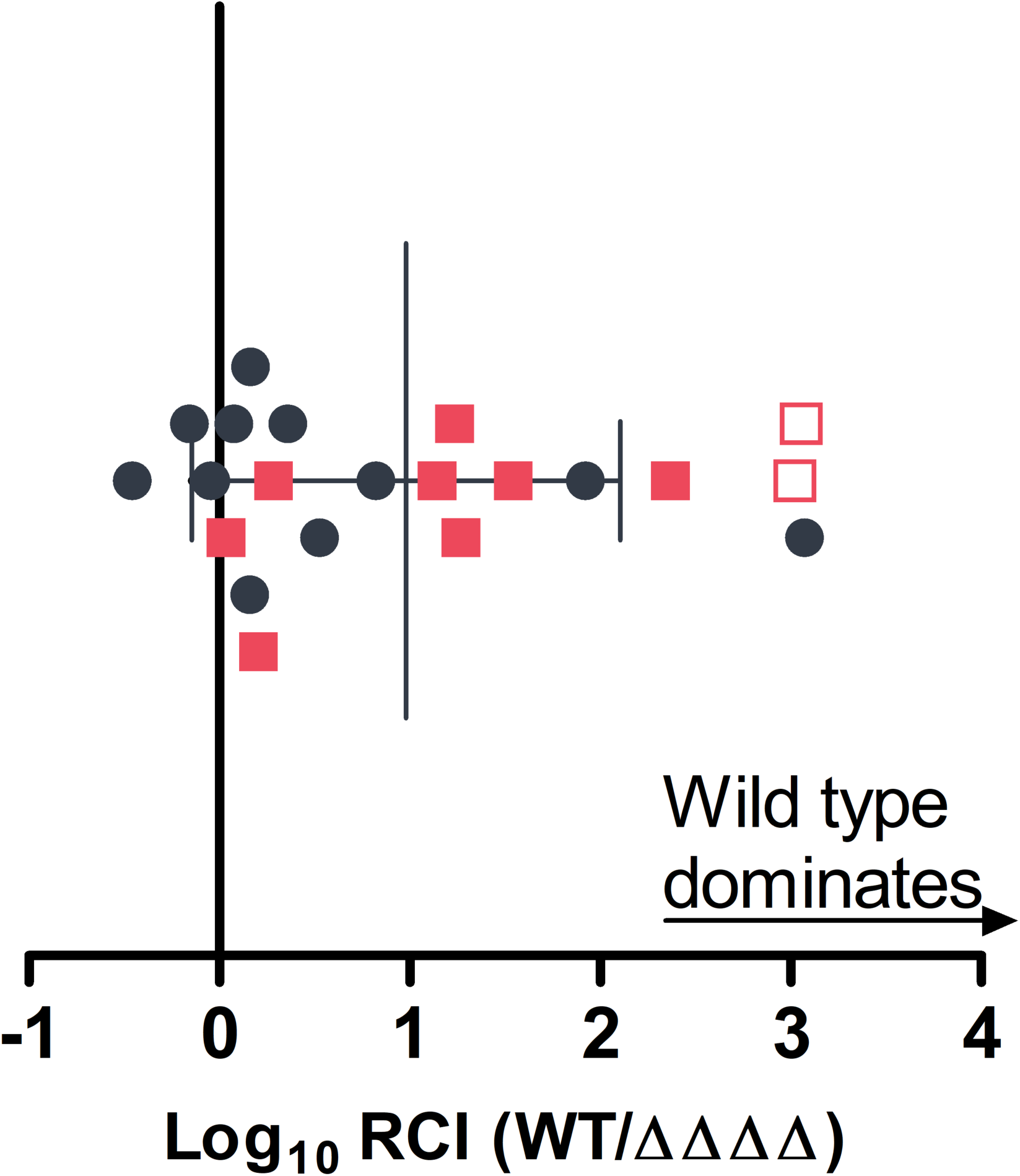
Light-organ colonization competition between wild-type *V. fischeri* and a chemotaxis mutant. Ratios of wild-type *V. fischeri* (WT) carrying a GFP-encoding plasmid (pVSV102) vs. unlabeled ΔVF_1133 ΔVF_2042 ΔVF_A0246 ΔVF_A0389 (“ΔΔΔΔ”) (circles) or unlabeled wild-type *V. fischeri* vs. ΔΔΔΔ carrying pVSV102 (squares) colonizing juvenile *E. scolopes* light organs were measured after 3 h of exposure to the inoculum followed by 21 h of incubation in sterile seawater. RCI, relative competitive index of bacterial strains. Each point represents one animal. Empty symbols represent the limit of detection for light organs in which only the wild-type strain was found. Error bars represent ± one standard deviation.

To determine whether the colonization defect displayed by ΔΔΔΔ is due to the chemosensory function of any of the four MCP genes, we sought to identify the chemoattractants recognized by these MCPs. Wild-type *V. fischeri* and MCP deletion mutants were spotted onto minimal-medium, soft-agar plates containing 1 mM of three known chemoattractants for *V. fischeri* (12): *N-*acetyl-D-glucosamine (GlcNAc), *N,N*′-diacetylchitobiose ((GlcNAc)_2_), or *N*-acetylneuraminic acid (Neu5Ac). The chemotactic zone sizes for ΔVF_1133 and ΔVF_A0246 were significantly smaller than for wild-type *V. fischeri* on each of the *N*-acetylated sugars (Table 3). Oddly, while the zone sizes of ΔΔΔΔ on GlcNAc and (GlcNAc)_2_ were similar to those of ΔVF_1133 and ΔVF_A0246, on Neu5Ac the ΔΔΔΔ zone size was comparable to wild type. This is a repeatable phenotype we have yet to explain. However, a ΔVF_1133 ΔVF_A0246 mutant produced a similar zone size to the single mutants on Neu5Ac. There was no significant difference in zone sizes between ΔΔΔΔ and wild-type *V. fischeri* spotted onto plates containing either no chemoattractant or 1 mM glucose (Table 3), indicating these mutations do not affect motility or chemotaxis in general. Because they grew equally well on either GlcNAc or Neu5AC (Fig. S2), the differences in zone sizes of the MCP mutants and wild-type *V. fischeri* were unlikely to be growth-related.

**Table 3.**
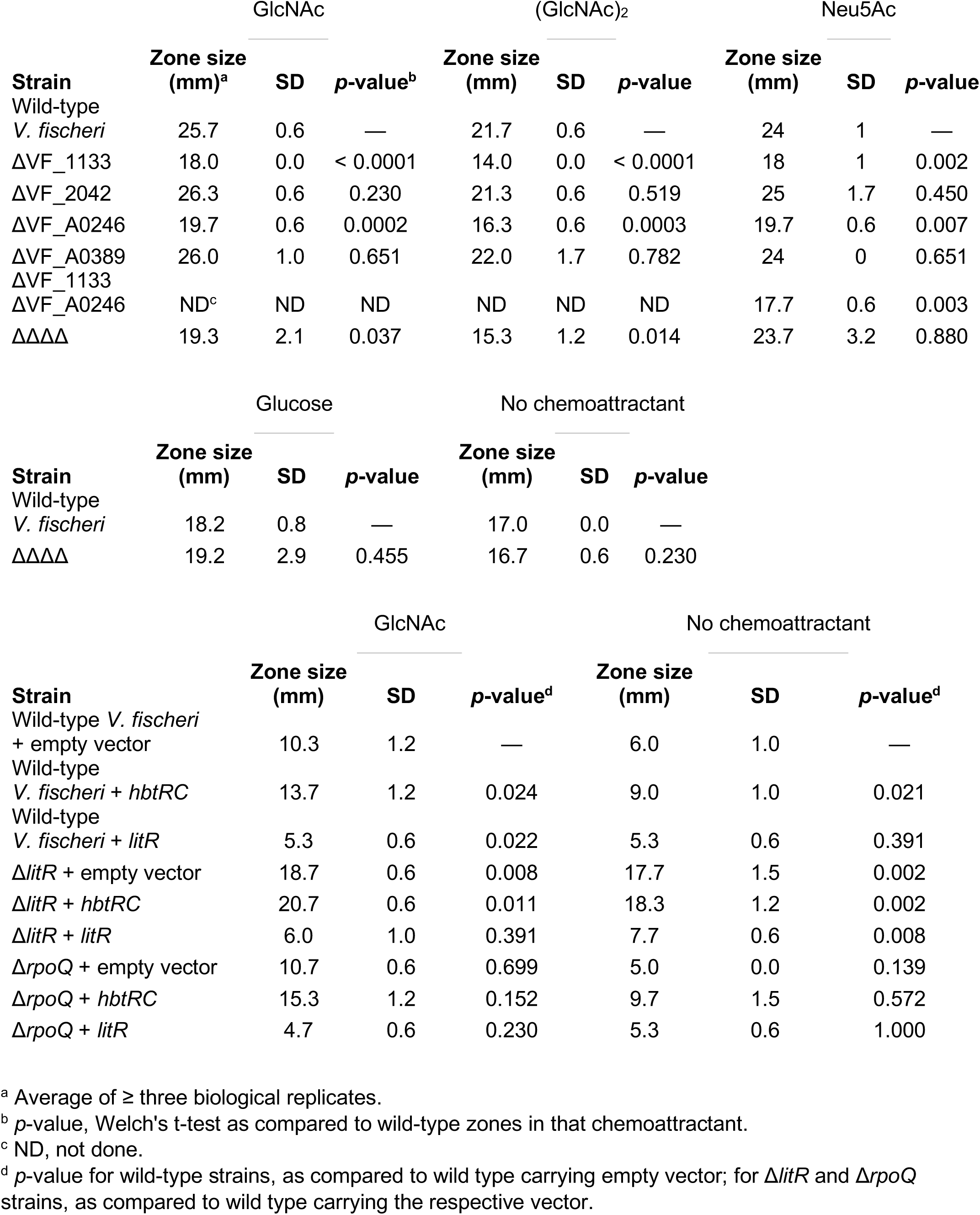
Soft-agar migration zone sizes.

### HbtR activates and LitR represses motility

Two flagellin genes, *flaC* and *flaF*, were upregulated by *hbtRC* (Table 1, Table S1), indicating the possible activation of motility by HbtR. An earlier study demonstrated that LitR represses motility in *V. fischeri* (31), and microarrays performed previously (Studer, Schaefer, and Ruby; Schaefer and Ruby unpublished data) indicated that AinS and LitR downregulate the expression of a majority of the flagellin genes within the locus VF_1836–77, as well as *flaF, fliL2, motX, motA1B1*, and *cheW*. To confirm the repression of motility genes by LitR, RT-qPCR was employed to determine the level of expression of three flagellin or motor genes in different genomic regions in wild-type *V. fischeri* and Δ*litR*. Expression of *cheW, motA1, flaC*, and *flaF* was moderately, but statistically significantly, higher in Δ*litR* than in wild-type *V. fischeri* (Table 2). However, we were not able to determine the mechanism by which LitR represses multiple motility gene loci, as expression levels of the known flagellin regulatory genes *rpoN, flrA, flrB, fliA*, and *flgM* were comparable between Δ*litR* and wild-type *V. fischeri* strains (Table 2).

The sigma factor-like regulator RpoQ, which is upregulated by LitR, represses motility and/or chemotaxis towards GlcNAc when overexpressed (32). Thus, we wanted to determine whether the effect LitR has on motility and *N*-acetylated sugar chemotaxis is mediated through its regulation of *rpoQ*. On soft-agar chemotaxis plates containing no chemoattractant or 1 mM GlcNAc, we observed the same pattern regardless of the presence of GlcNAc: either deletion of *litR* or overexpression of *hbtRC* led to larger migration zones than for wild-type *V. fischeri*, while overexpression of *litR* essentially eliminated bacterial migration (Table 3). There were no significant differences between Δ*rpoQ* and wild-type *V. fischeri* strains carrying the same vector (Table 3), indicating that the repression of motility by LitR is not mediated through RpoQ.

### HbtR represses and LitR activates exopolysaccharide production

An approximately 25-kb locus constituting the genes VF_0157–80 was downregulated when *hbtRC* was expressed (Table 1, Table S1). Previously performed microarrays indicated that most of the genes in this locus (VF_0159 and VF_0162–0180) are also upregulated by AinS and LitR (Studer, Schaefer, and Ruby; Schaefer and Ruby, unpublished data). To confirm the upregulation of this locus by LitR, RT-qPCR was used to evaluate the expression of three genes, located in separate operons, in Δ*litR* and wild-type *V. fischeri*. Expression of VF_0157 (*wbfB*), VF_0164 (*etp*), and VF_0165 (*wzc*) was significantly higher in wild-type *V. fischeri* than in Δ*litR* (Table 2). Expression of *etp* was significantly lower in Δ*hbtRC* constitutively expressing *hbtRC* compared to Δ*hbtRC* carrying empty vector (Table 2), confirming that HbtR regulation of *litR* represses gene expression in this locus.

Most of the genes in the locus VF_0157–80 are annotated as being involved in production of extracellular polysaccharides. However, gene annotation alone could not indicate which component(s) of the cell envelope (lipopolysaccharide O-antigen, capsule, or exopolysaccharide (EPS)) might be affected by the genes in this locus. To determine which polysaccharide type is produced by the enzymes encoded in VF_0157–80, alcian blue staining was used to detect negatively charged polysaccharides in the supernatants of liquid cultures of *V. fischeri* strains. Both Δ*litR* and ΔVF0157–80 produced less alcian blue-staining EPS than wild-type *V. fischeri*, with ΔVF_0157–80 producing even less EPS than Δ*litR* (Fig. 3). Overexpression of *hbtRC* repressed EPS production in wild-type *V. fischeri*, but not in Δ*litR* or ΔVF_0157–80 (Fig. 3). Overexpression of *litR* in ΔVF_0157–80 did not complement the reduction in EPS produced by that mutant (Fig. 3), indicating that essentially all EPS is produced by this locus.

**FIG 3.**
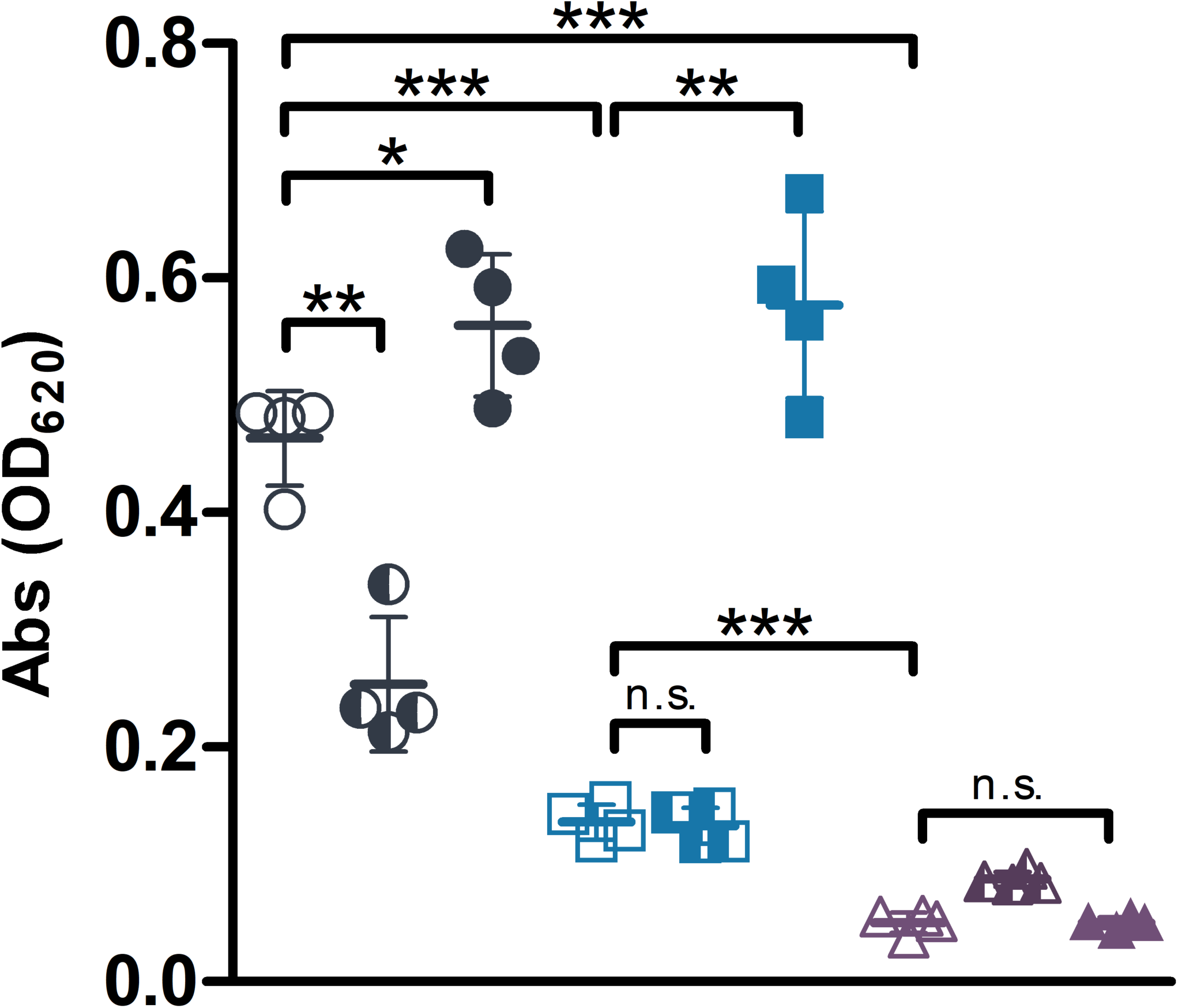
EPS production in wild-type *V. fischeri*, Δ*litR*, and ΔVF_0157–80 strains. EPS in supernatants of 24-h cultures grown in 0.05% casamino acids and 6.5 mM Neu5Ac was stained with alcian blue for wild-type *V. fischeri* (circles), Δ*litR* (squares), and ΔVF_0157–80 (triangles) carrying empty pVSV105 (open symbols), pVSV105::*hbtRC* (half-filled symbols), or pVSV105::*litR* (closed symbols). Error bars represent ± one standard deviation. Abs, absorbance.*, *p*<0.05; **, *p*<0.005; ***, *p*<0.001; n.s., not statistically significant.

### *hbtRC* expression is regulated by Crp, but not by AphB or quorum sensing

To determine whether *hbtRC* expression in *V. fischeri* is affected by quorum sensing and/or Crp, a regulator known to affect *tcpPH* expression in *V. cholerae*, RT-qPCR was performed on wild-type *V. fischeri*, Δ*litR*, and Δ*crp* grown to OD_600_ ∼0.3 and ∼1.0. Expression of *hbtRC* was lower at high cell density than at low cell density for all three strains, with no difference between wild-type *V. fischeri* and Δ*litR* at either higher or lower OD_600_ (Table 2). Expression of *hbtRC* in Δ*crp* was below that in wild-type *V. fischeri* at low OD_600_, but comparable at high OD_600_. Thus, Crp appears to activate *hbtRC* expression, the reverse of *tcpPH* regulation by Crp in *V. cholerae*.

Because Crp activates quorum-signaling genes in the *V. fischeri* genome (33, 34), we asked whether quorum signaling is responsible for the change in *hbtRC* expression at different culture densities, and whether this regulation explains the lower expression in the Δ*crp* mutant. The RT-qPCR experiment was repeated for wild-type *V. fischeri* grown in SWT with or without added autoinducers. *hbtRC* expression remained higher at lower cell density in the presence of both autoinducers (Table 2), indicating that quorum signaling is not involved in Crp- and cell density-mediated *hbtRC* regulation.

To determine whether the regulator AphAB and/or a change in pH affect *hbtRC* expression in *V. fischeri*, RT-qPCR was performed on RNA extracted from cultures of wild type or Δ*aphB V. fischeri* grown at pH 5.5 or 8.5. There was no significant difference in *hbtRC* expression between any of the conditions (Table S2).

### *tcpPH* homologs do not cross-complement between *V. fischeri* and *V. cholerae*

Δ*tcpPH* and Δ*hbtRC* strains of *V. cholerae* and *V. fischeri*, respectively, were complemented with empty vector, vector expressing *tcpPH*, or vector expressing *hbtRC*. RT-qPCR was performed on RNA extracted from cultures of each strain to determine the level of *toxT* expression in the *V. cholerae* strains or *litR* expression in the *V. fischeri* strains. While expression of each *tcpPH* homolog complemented the respective deletion, cross-complementation had no effect on output gene expression (Table S2), illustrating the divergent activities of HbtR and TcpP.

In *V. cholerae*, ToxR cooperates with TcpP to activate *toxT* expression (35, 36), begging the question of whether ToxR is also involved in controlling the *V. fischeri* HbtR regulon. RT-qPCR performed on RNA extracted from cultures of Δ*hbtRC* and Δ*hbtRC*Δ*toxRS* carrying either empty vector or vector expressing *hbtRC* showed that *litR* expression was repressed to the same degree by HbtR regardless of the presence or absence of *toxRS* (Table S2), further demonstrating that regulation by HbtR is independent of ToxR and differs substantially from the TcpP system.

### Δ*hbtRC* has no defect in colonizing juvenile *E. scolopes* light organs

Previous work indicated that Δ*hbtRC* (then referred to as Δ*tcpPH*) had a light-organ colonization defect, which appeared to increase through 96 h (19). We replicated this defect when using the same conditions as the previous study (Fig. 4A); however, we observed an advantage for Δ*hbtRC* when we reversed the fluorescent marker plasmids (Fig. 4A), indicating the earlier results were due simply to experimental design. Co-colonization of juvenile *E. scolopes* with chromosomally labeled strains resulted in no significant colonization defect for either strain (Fig. 4B), demonstrating that HbtR is, in fact, not required for entry into, or life inside, the *E. scolopes* light organ. Based on this experience, we urge caution by other researchers using *rfp*-labeled pVSV208 in co-colonization experiments.

**FIG 4.**
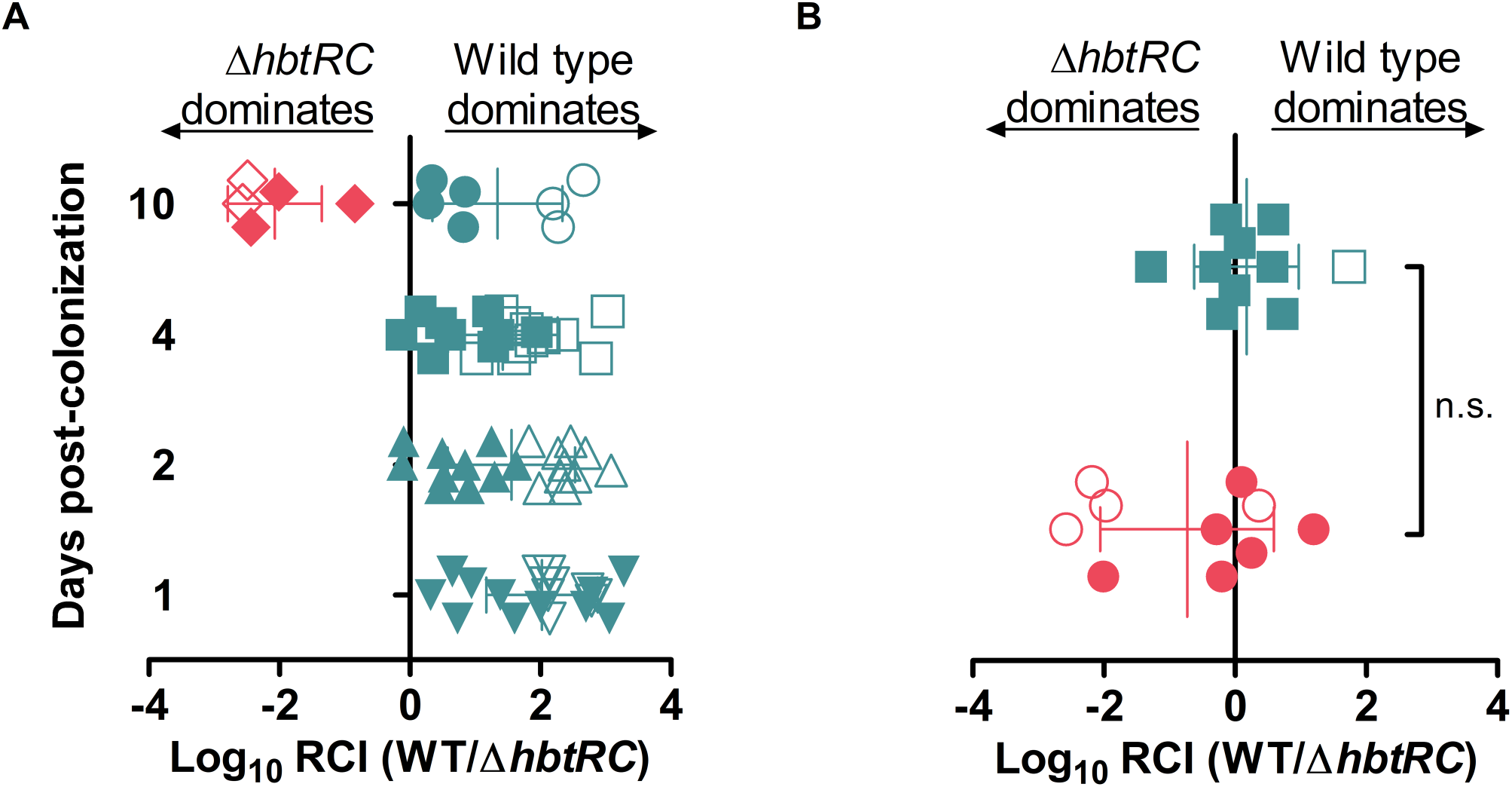
Light-organ colonization competition between wild-type *V. fischeri* and Δ*hbtRC* strains. (A) Ratios of wild-type *V. fischeri* carrying a GFP-encoding plasmid (pVSV102) vs. Δ*hbtRC* carrying an RFP-encoding plasmid (pVSV208) (circles, squares, and triangles) or wild-type *V. fischeri* (WT) carrying pVSV208 vs. Δ*hbtRC* carrying pVSV102 (diamonds) colonizing juvenile *E. scolopes* light organs were measured after 1 to 10 d. (B) Ratios of wild-type *V. fischeri* with chromosomal *gfp* vs. Δ*hbtRC* with chromosomal *rfp* (squares) or wild-type *V. fischeri* with chromosomal *rfp* vs. Δ*hbtRC* with chromosomal *gfp* (circles) colonizing juvenile *E. scolopes* light organs were measured after 1 d. RCI, relative competitive index of bacterial strains. Each point represents one animal. Empty symbols represent the limit of detection for light organs in which only the *gfp*-carrying strain was found. Error bars represent ± one standard deviation. n.s., not statistically significant.

### Transcription of *hbtR* is activated as symbionts exit the light organ

Considering the effects HbtR has on chemotaxis and luminescence gene expression (Tables 1, 2, and S1), and that *hbtRC* has no effect on squid luminescence (Fig. S1) or colonization (Fig. 4B), we postulated that HbtR was likely to be involved in the transition into planktonic life as *V. fischeri* exits the light organ. To establish at which point in the juvenile squid diel cycle *V. fischeri hbtRC* expression is activated, we performed *in situ* hybridization chain reaction (HCR) targeting *hbtR* mRNA to determine whether this gene was expressed in the bacteria before, during, or after their expulsion from the squid light organ into seawater. *hbtR* mRNA was not detected in *V. fischeri* cells within the crypts or during expulsion along the path out of the light organ (Fig. 5A). Only in those cells that had completely exited the light-organ pores was *hbtR* expression detected (Fig. 5B and Movie S1). Within an hour of expulsion into seawater, ∼25% of expelled *V. fischeri* cells were expressing *hbtR* (Fig. 5C). As expected, transcripts of the quorum signaling-induced gene *litR* were detected within *V. fischeri* cells inside the *E. scolopes* light-organ crypts (Fig. S3); thus, it is unlikely that HCR was unable to detect *hbtR* transcripts in colonizing bacteria due to failure of the reagents to penetrate the light organ.

**FIG 5.**
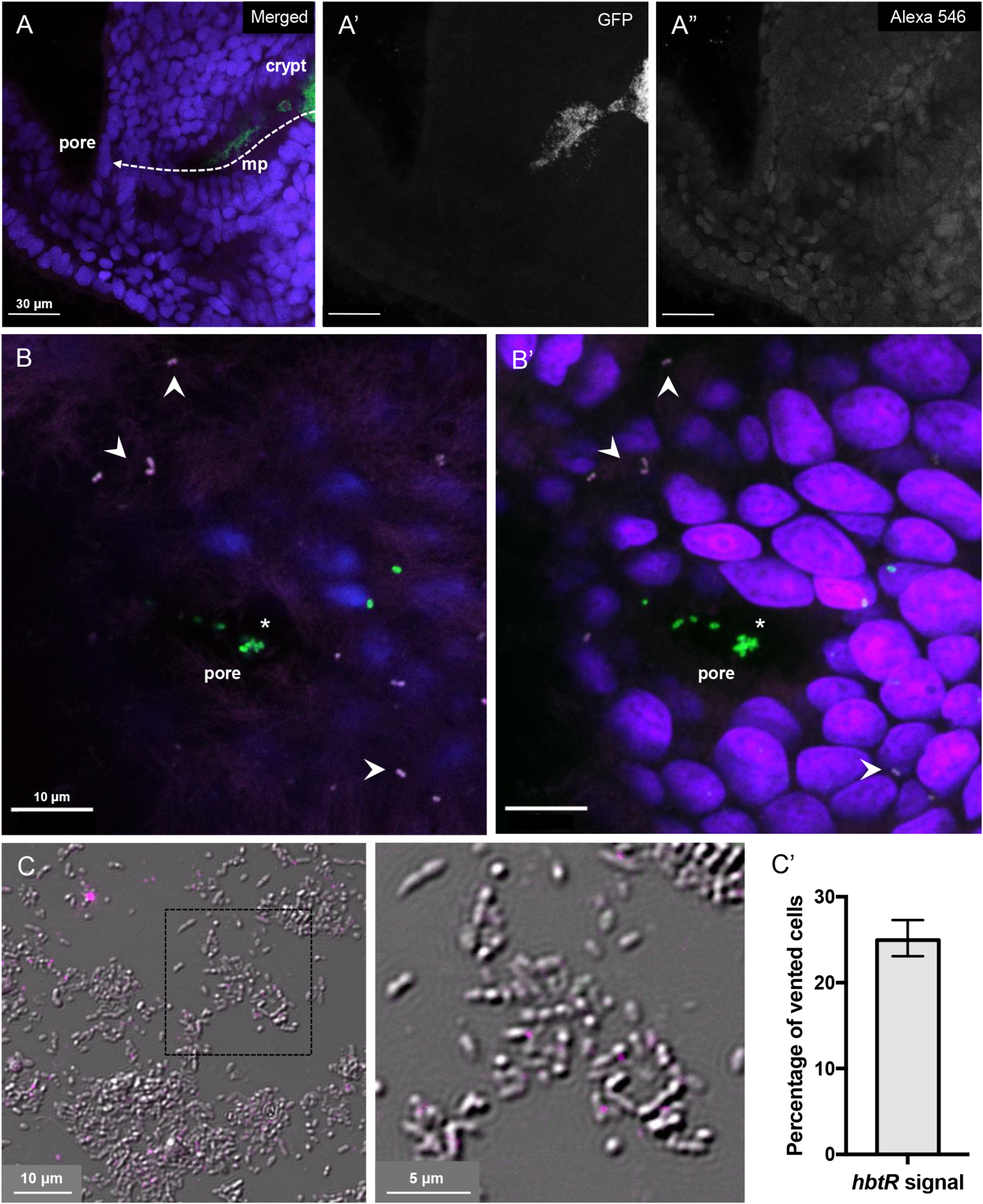
Expression of *hbtR* in *V. fischeri* during expulsion from *E. scolopes* light organs. Transcripts of *hbtR* (magenta, arrowheads) were detected using *in situ* HCR in *V. fischeri* cells (green) during expulsion from *E. scolopes* juvenile light organ crypts. (A) *V. fischeri* symbionts originate in the crypt, pass through the migration path (mp), and exit through the external surface pore; (A) merged image, (A’) GFP channel, (A”) *hbtR* transcript channel (Alexa 546). (B) *V. fischeri* cells (green) are expelled through the external surface pore (asterisk) into seawater 1 h after a dawn light cue; (B) a single optical slice of the superficial layer above the exit point from host and *V. fischeri* cells outside of the pore expressing *hbtR*; (B’), the corresponding merged composite stack (video in Movie S1). (C) One hour after expulsion into ambient seawater, vented cells express *hbtR* (magenta); enlargement is of the area within the dashed square. (C’) Percentage and 95% confidence intervals of vented cells expressing *hbtR* calculated from four samples of ventate on slides, from two experiments. Blue, *E. scolopes* nuclei (DAPI).

## Discussion

While much research has gone into understanding the mechanisms necessary for colonization of the *E. scolopes* light organ by *V. fischeri* (reviewed most recently in (37)), there has been little investigation into how the ∼95% of symbiotic bacteria expelled from the light organ at dawn each day transition back into life in seawater. In this work we present HbtR, a heterofunctional homolog of the *V. cholerae* virulence regulator TcpP that represses the quorum-signaling regulator LitR. While it is unclear at this time whether HbtR represses *litR* expression directly or through some intermediate regulator(s), the RNA-seq data indicate HbtR does not act through a step in the quorum-signaling pathway upstream of LitR (Table S1). Through the discovery of additional functions regulated by LitR, namely chemotaxis, motility, and EPS production, we have begun to build a picture in which LitR aids the transition by *V. fischeri* into a colonization lifestyle and, upon expulsion from the light organ, HbtR reverses that process to help transition back into planktonic life (Fig. 6). This is a markedly different role for HbtR from that of TcpP.

**FIG 6.**
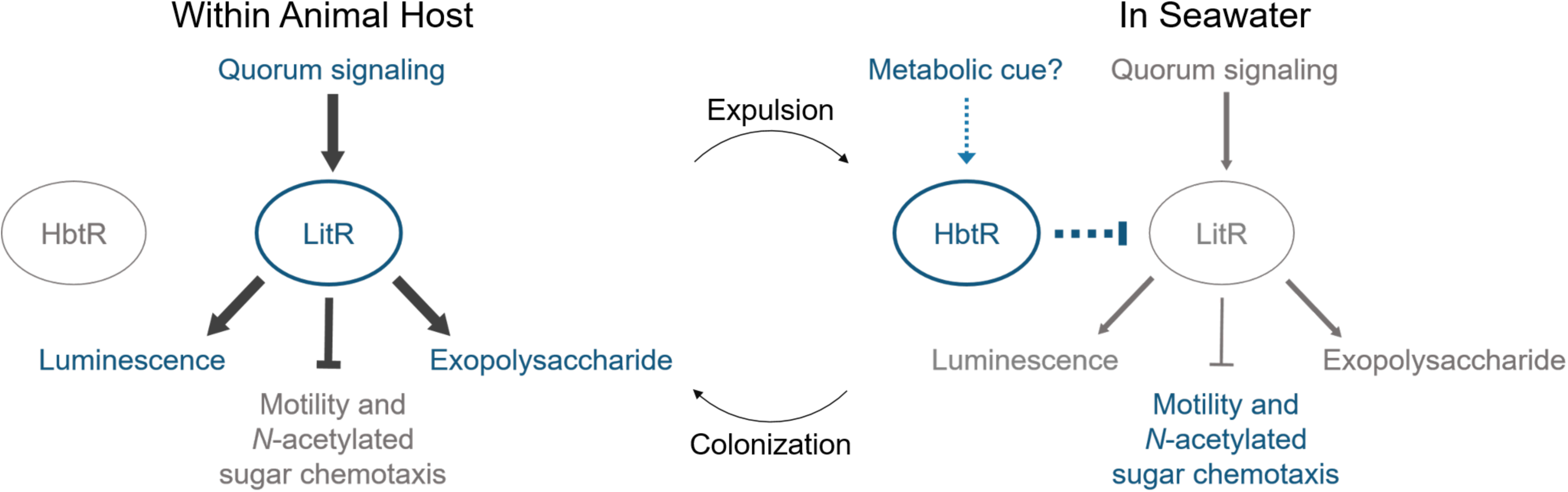
Model proposing regulation by HbtR and LitR in *V. fischeri* during colonization of and expulsion from the *E. scolopes* light organ. Upon colonization, LitR regulates relevant physiological phenotypes, including activation of luminescence and EPS production and repression of motility and chemotaxis. HbtR causes the repression of *litR* expression upon expulsion of *V. fischeri* from the light organ, reversing this process and leading to the upregulation of motility and chemotaxis and the downregulation of luminescence and EPS production. Gray text, downregulated genes and phenotypes; blue text, upregulated genes and phenotypes.

The regulatory mechanisms controlling *hbtRC* expression, as elucidated thus far, are similar to the *tcpPH* regulatory pathway in *V. cholerae* only in that both operons are regulated by the global regulator Crp, albeit in the opposite manner. Notably, the *hapR* homolog in *V. fischeri, litR*, is itself repressed by HbtR (Tables 1, S1, S2), a reversal of roles from those in *V. cholerae*. Considering the distinct regulatory pathways for HbtR and TcpP, as well as the low percent identity (26.8%) of the two protein sequences, it appears, evolutionarily, that either progenitors of *tcpPH* and *hbtRC* were acquired independently after separation of the two species’ ancestors, or the parent operon evolved in dramatically different fashions as the two species evolved to inhabit disparate environmental niches. Determining which of these histories is the more likely is beyond the scope of this study, but in either case it is clear the two homologs have been recruited into specialized roles befitting the two species’ distinct lifestyles, i.e., pathogenesis versus beneficial symbiosis within animal hosts.

The data presented here suggest HbtR acts as a first-wave responder as *V. fischeri* bacteria exit the light organ and re-enter the seawater (Fig. 5), a significant change in environmental conditions and an imperative for survival. This role could explain some of the discrepancies observed between the RNA-seq results in the present study and those in Thompson 2017 (19). For example, the Thompson data indicate that in addition to *hbtR* and *hbtC*, the luminescence genes *luxR, luxI*, and *luxCDABEG* were also highly upregulated in cells recently expelled from the light organs compared to planktonic cells, while *litR*, which activates the *lux* genes, was not differentially regulated between the two cell populations. Meanwhile, the results of this study indicate that HbtR represses LitR and consequently causes the downregulation of *lux* and other downstream quorum-signaling genes (Tables 1, S1, S2). Similarly, the gene most highly upregulated by HbtR in the RNA-seq data presented here was VF_2042 (Table S1), one of the four MCP genes repressed by LitR (Table 2); in the earlier study, VF_2042 was downregulated in cells released from the light organs (19). These results are consistent with a model in which HbtR becomes active once *V. fischeri* exits the light organ, with some of the earliest effects seen in its repression of *litR*. In the earlier study, it may have been that in the ∼25 minutes between initiation of light-organ expulsion and harvesting the *V. fischeri* RNA, HbtR had become activated (Fig. 5) and was already repressing *litR* expression, but the downstream effects (e.g., on VF_2042 and *lux* gene expression) were not yet apparent.

Previous studies have also demonstrated a repression of several flagellin genes by AinS and increased motility in a *litR* mutant (31). Here we show that LitR represses numerous flagellin, motor, and chemotaxis genes, including at least two MCP genes involved in chemotaxis toward (GlcNAc)_2_, an *N*-acetylated sugar reported to be an important chemoattractant during light-organ colonization (12). Previous work indicated that a *litR* mutant colonized *E. scolopes* to a lower level than wild-type *V. fischeri* (31), which the authors suggested was due to an initiation delay due to hypermotility. However, in the study in which LitR was first described (4), a *litR* mutant had an advantage in initiating light-organ colonization over wild-type *V. fischeri.* Based on work presented here, including a demonstration that MCP genes repressed by LitR are involved in light-organ colonization (Fig. 2), we posit that the *litR-*mutant colonization benefit (4) may have resulted from the increase in both motility and chemotaxis into the light organ by that mutant. We hypothesize that the lower level of colonization observed for the *litR* mutant in single-strain experiments (31) may have been due to dysregulation of a separate symbiotic process, perhaps extracellular polysaccharide production.

LitR appears to activate, and HbtR to thus repress, a large locus of genes (VF_0157–80) involved in extracellular polysaccharide production (Tables 1, S1, S2). Fidopiastis et al. (4) noted that *litR*^-^ colonies are less opaque than wild-type *V. fischeri* and suggested this phenotype may be due to a difference in the extracellular polysaccharides present in each strain. The nature of polysaccharide affected by VF_0157–80 is not fully established, yet it seems most likely to be an EPS, or “slime layer,” due to the ease with which it separates from the bacterial cell during centrifugation (Fig. 3A). While lipopolysaccharide and the *syp* polysaccharide have been implicated in the initiation of host colonization by *V. fischeri* (38–40), little has been determined regarding the possible involvement of extracellular polysaccharides in colonization persistence. *V. fischeri*-colonized crypts, despite having no goblet cells (Nyholm, personal communication), have been shown to stain alcian blue-positive (41), indicating that the presumptive EPS made by VF_0157–80 may be a relevant factor in *E. scolopes*-*V. fischeri* symbiosis. This aspect of the symbiotic relationship could have broader implications for host-microbe interactions in general.

## Materials and Methods

### Bacterial strains and growth conditions

Bacterial strains and plasmids used in this study are summarized in Table S3. *E. coli* and *V. cholerae* were grown in lysogeny broth (LB; (42), (19)) at 37°C; *V. fischeri* was grown in LB-salt (LBS; (43)) at 28°C. Overnight cultures were inoculated with single colonies from freshly streaked −80°C stocks. Liquid cultures were shaken at 225 rpm. Unless otherwise noted, 15 g of agar were added per liter for solid media. Antibiotics were added to overnight cultures, where applicable, at the following concentrations: kanamycin (Km), 50 μg/mL; chloramphenicol (Cm), 5 μg/mL (*V. fischeri*) or 25 μg/mL (*E. coli*). Experimental cultures were grown in either seawater-tryptone (SWT) medium (44) or minimal-salts medium (MSM; per L: 8.8 g NaCl, 6.2 g MgSO_4_·7H_2_O, 0.74 g CaCl_2_·2H_2_O, 0.0204 g H_2_NaPO_4_, 0.37 g KCl, 8.66 g PIPES disodium salt, 0.65 mg FeSO_4_·7H_2_O; pH 7.5) supplemented with 5 mg/L casamino acids and a carbon source as indicated. The results of all experiments are reported as the means of three (or more, as indicated) biological replicates ± 1 standard deviation (SD). Statistical analysis was performed using Welch’s t-test.

### Plasmid and mutant construction

Primers used to construct the deletion and expression vectors in this study are listed in Table S4. In-frame deletion of genes from the *V. fischeri* genome was performed as previously described (45, 46). Counter-selection to remove the target gene and pSMV3 was performed on LB-sucrose (per L: 2.5 g NaCl, 10 g Bacto Tryptone, 5 g yeast extract, 100 g sucrose) for two days at room temperature. Expression vectors were constructed by amplifying and ligating the target gene into the multiple cloning site of pVSV105 (47). Inducible expression of *hbtRC* was achieved by ligating *lacI*^*q*^ from pAKD601 to *hbtRC* cloned from the *V. fischeri* genome and inserting the fusion into the multiple cloning site of pVSV105. Overexpression of *hbtRC* brought the level of expression closer to that found upon expulsion from juvenile squid (19), which was higher than we measured for wild-type *V. fischeri* in culture conditions. Genomic insertion of *gfp* or *rfp* under control of the *lac* promoter at the *attTn7* site was performed using a mini-Tn7 vector as described (48). Previously constructed Δ*hbtRC* (19) was used to recapitulate conditions for follow-up on previous experiments (RNA-seq and squid co-colonization); a newly derived Δ*hbtRC* strain was used for all other experiments so it would have the same parent wild-type *V. fischeri* stock as other mutants created in this study.

### Generation and analysis of RNAseq libraries

RNA extraction and RNAseq were performed as previously described (19). Briefly, RNA was stabilized with RNAprotect Bacterial Reagent (Qiagen) and extracted with an RNeasy Mini Kit (Qiagen) from 500 μL of cultures grown to mid-log phase in SWT with 1.75 mM IPTG. RNA was extracted from biological triplicates for each strain. Contaminating DNA was degraded by treatment with TURBO DNase (Invitrogen). Ribosomal reduction, strand-specific library preparation, and paired-end 50-bp sequence analysis on an Illumina HiSeq 2500 on high-output mode were performed at the University of Minnesota Genomics Center. Sequences were processed on the open-source Galaxy server (usegalaxy.org; (49)) using the following workflow (default settings used unless otherwise indicated): reads were trimmed with Trimmomatic (maximum mismatch:1, LEADING:3, TRAILING:3, SLIDINGWINDOW:4,20, MINLEN:35), mapped to the *V. fischeri* chromosomes and plasmid (accession nos. NC_006840.2, NC_006841.2, and NC_006842.2) using Bowtie2 (-un-con, --sensitive), and counted with featureCounts (-p disabled, -t=gene, -g=locus_tag, -Q=10, -s); differential expression was analyzed with DESeq2 (outliers filtering and independent filtering turned on). Between 11.7 million and 12.7 million paired reads were mapped to the *V. fischeri* genome per biological replicate. Raw and processed read files have been deposited into the NCBI Gene Expression Omnibus server (GSE151621).

### Reverse transcription and quantitative PCR

RNA from bacterial cultures was extracted as described above and treated with TURBO DNase (Invitrogen). cDNA was synthesized from DNase-treated RNA using SMART MMLV Reverse Transcriptase (Clontech) and Random Hexamer Primer (Thermo Scientific). Gene expression was measured by qPCR performed with LightCycler 480 SYBR Green I Master Mix (Roche) under the following conditions: 95°C for 5 min; 45 cycles of 95°C for 10 sec, 60°C for 20 sec, and 72°C for 20 sec; melting curve acquisition from 65°C to 97°C. qPCR primer pairs were designed for these conditions and confirmed to have efficiencies between 90 and 110%. Cycle thresholds (Ct) for each sample were normalized to those of the reference gene *polA* (ΔCt = target gene-*polA*; *polA* expression primers provided courtesy of Silvia Moriano-Gutierrez). ΔΔCt values were calculated by subtracting the average ΔCt for the parent strain, at higher pH or lower OD_600_ when appropriate; fold changes were calculated as 2^(-ΔΔCt)^.

### Growth and luminescence curves

Overnight LBS cultures of each strain were pelleted, washed once, and resuspended in 1 mL of the intended growth medium. Luminescence growth curves were performed in 20 mL SWT in flasks shaking at 225 rpm; periodic 1 mL and 300 μL aliquots were taken for luminescence and OD_600_ readings on a luminometer (Turner Designs) and Tecan Genios plate reader, respectively. Growth curves were performed in 1.2 mL growth medium and monitored in the Tecan Genios plate reader with continuous shaking at high speed.

### Squid colonization assays

Juvenile squid colonization experiments were performed as described previously (50), with juvenile squid exposed to *V. fischeri* strains at 1,000– 6,000 CFU/mL either for 3 h or overnight, as indicated, before being placed in fresh sterile seawater until euthanizing by freezing. Colonization competition experiments were performed by exposing juvenile squid to a 1:1 inoculum of each strain.

### *In situ* hybridization chain reaction

HCR (51) was performed on squid colonized with wild-type *V. fischeri* carrying a *gfp*-labeled plasmid and on *V. fischeri* cells 1 h after being expelled from squid light organs as previously described (52–54). Briefly, *E. scolopes* juveniles and expelled *V. fischeri* cells were fixed in 4% paraformaldehyde in marine phosphate-buffered saline either before or after a light cue stimulated bacterial expulsion. HCR was performed on dissected light organs (52) and expelled *V. fischeri* cells affixed to gelatin-coated slides (adapted from (53)) using HCR version 3.0 chemistry (54). Ten probes targeting *hbtR* mRNA and eleven probes targeting *litR* mRNA (Table 3) were amplified with Alexa Fluor 546-labeled hairpins (Molecular Instruments). Light organs were then counterstained overnight with a 1:750 dilution of 4′,6-diamidino-2-phenylindole (DAPI; Thermo Fisher Scientific) in 5x SSC-Tween before mounting on slides with Vectashield (Vector Laboratories) and overlaid with a coverslip (#1.5, Fisherbrand, Fisher Scientific). Imaging was done on an upright Zeiss LSM 710 laser-scanning confocal microscope at the University of Hawai‘i Kewalo Marine Laboratory; images were analyzed using FIJI (ImageJ) (55).

### Motility and chemotaxis assays

Soft-agar motility assays were performed as previously described (56). Briefly, the equivalent of 10 μL of a bacterial culture grown to an OD_600_ of 0.5 in SWT were spotted onto minimal-medium plates containing 0.25% agar and, when appropriate, 1 mM chemoattractant. Migration zones were measured after 18–24 h of static incubation at 28°C.

### Alcian blue detection of extracellular polysaccharides

EPS was detected by staining with alcian blue essentially as previously described (57). Single colonies from freshly streaked −80°C stocks were used to inoculate MSM supplemented with 6.5 mM Neu5Ac and 0.05% (wt/vol) casamino acids and grown with shaking 18–22 h. EPS was separated from cells by centrifugation of the equivalent of 2.5 mL at OD_600_=1.0 for 15 min at 12,000 xg and 4°C. 250 μL of this supernatant was mixed with 1 mL alcian-blue solution (per L: 0.5 g alcian blue, 30 mL glacial acetic acid, pH 2.5) and rocked for 1 h at room temperature, then centrifuged for 10 min at 10,000 rpm and 4°C. Pellets were resuspended in 1 mL 100% ethanol, then centrifuged for 10 min at 10,000 xg and 4°C. Pellets were solubilized in 500 μL SDS (per L: 100 g sodium dodecyl sulfate, 50 mM sodium acetate, pH 5.8), and the absorbance was read at OD_620_.

## Acknowledgments

The authors thank Silvia Moriano-Gutierrez and Alexandrea Duscher for advice regarding RNA-seq, Elliott McCloskey and Roxane Gaedeke for help with experiment execution, the University of Minnesota Genomics Center for RNA processing and next-generation sequencing, and the Ruby and McFall-Ngai laboratories for their insights. The research was supported by NIH grants R37 AI50661 (Margaret McFall-Ngai and E.G.R.), R01 OD11024, and R01 GM135254 (E.G.R. and M.M.-N.) and NSF INSPIRE Grant MCB1608744 (Eva Kanso and Scott Fraser, University of Southern California). Acquisition of the Leica TCS SP8 X confocal was supported by NSF DBI 1828262 (Marilyn Dunlap, E.G.R., and M.M.-N.), and microscopy was performed in the MICRO facility, supported by COBRE P20 GM125508 (M.M.-N. and E.G.R.).

## SUPPLEMENTAL MATERIAL LEGENDS

**FIG S1** Luminescence of wild-type *V. fischeri* and Δ*hbtRC* strains during symbiosis. Luminescence was measured in juvenile squid 24 h or 48 h after colonization with either wild-type *V. fischeri* (●) or Δ*hbtRC* (■). Each point represents one animal. Mean values indicated; no significant differences were noted between strains at either time-point. RLU, relative light units.

**FIG S2** Growth of wild-type *V. fischeri* and chemotaxis mutants on *N*-acetylated sugars. (A) The rates of growth in MSM supplemented with 0.05% casamino acids and 6.5 mM Neu5Ac were measured for wild-type *V. fischeri* (●), ΔΔΔΔ (■), and ΔVF_1133 ΔVF_A0246 (▴). (B) The rates of growth in MSM supplemented with 0.05% casamino acids and 6.5 mM GlcNAc were measured for wild-type *V. fischeri* (●) and ΔΔΔΔ (■). Results represent means of three biological replicates ± one standard deviation. Abs, absorbance.

**FIG S3** Expression of *litR* in *V. fischeri* when host-associated within the *E. scolopes* light organ. Transcripts of *litR* (green) were detected using *in situ* HCR in *V. fischeri* cells colonizing *E. scolopes* crypts (C) after traversing the tissues of the migration path (MP). Blue, *E. scolopes* nuclei (DAPI); green, *litR* transcripts (Alexa 546). Bar 15 μm.

**TABLE S1 RNA-seq data file** Expression-level counts of all orfs in the *V. fischeri* genome. Comparison of mapped read counts for Δ*hbtRC* and *hbtRC* (induced)

**TABLE S2 RT-qPCR results**

**TABLE S3 Bacterial strains and plasmids used in this work**

**TABLE S4 Primers and probes used in this work**

**MOVIE S1** Expression of *hbtR* in *V. fischeri* cells during their expulsion from an *E. scolopes* light organ. *hbtR* transcripts were detected using *in situ* HCR in *V. fischeri* cells expelled from the external surface pore of the light organ 1 h after a dawn light cue. Merged stack of images depicted in Fig 5B’. Blue, *E. scolopes* nuclei (DAPI); green, *gfp*-labeled *V. fischeri*; magenta, *hbtR* transcripts (Alexa 546).

## Notes

### Competing Interest Statement

The authors have declared no competing interest.

