## Supplemental files for "HbtR, a heterofunctional homolog of the virulence regulator TcpP, facilitates the transition between symbiotic and planktonic lifestyles in *Vibrio fischeri*"

### Table S2. RT-qPCR results

| | Wild-type *V. fischeri* | | | | Δ*aphB* | | | |
|---|---|---|---|---|---|---|---|---|
| **pH**[a] | **ΔCt (*hbtRC-polA*)** | **SD** | ***p*-value**[b] | **Fold change**[b] | **ΔCt (*hbtRC-polA*)** | **SD** | ***p*-value** | **Fold change** |
| 8.5 | 10.94 | 0.37 | — | — | 10.42 | 0.33 | 0.167 | 1.46 |
| 5.5 | 10.55 | 0.14 | 0.227 | 1.32 | 10.34 | 0.34 | 0.132 | 1.54 |

| **Strain**[c] | **ΔCt (*litR-polA*)** | **SD** | ***p*-value** | **Fold change** |
|---|---|---|---|---|
| Δ*hbtRC* + empty vector | -4.38 | 0.19 | — | — |
| Δ*hbtRC* + *hbtRC* | -3.46 | 0.42 | 0.032 | 0.54 |
| Δ*hbtRC* + *tcpPH* | -4.65 | 0.05 | 0.131 | 1.20 |

| **Strain**[d] | **ΔCt (*toxT-polA*)** | **SD** | ***p*-value** | **Fold change** |
|---|---|---|---|---|
| Δ*tcpPH* + empty vector | 4.90 | 0.40 | — | — |
| Δ*tcpPH* + *hbtRC* | 4.61 | 0.52 | 0.498 | 1.27 |
| Δ*tcpPH* + *tcpPH* | -0.06 | 0.28 | 0.0004 | 31.65 |

| **Strain**[c] | **ΔCt (*litR-polA*)** | **SD** | ***p*-value** | **Fold change** |
|---|---|---|---|---|
| Δ*hbtRC* + empty vector | -2.19 | 0.17 | — | — |
| Δ*hbtRC* + *hbtRC* | -0.29 | 0.32 | 0.003 | 0.27 |
| Δ*hbtRC*Δ*toxRS* + empty vector | -1.57 | 0.96 | 0.188 | 1.19 |
| Δ*hbtRC*Δ*toxRS* + *hbtRC* | -0.72 | 0.26 | 0.004 | 0.37 |

[a] Cultures grown in MSM buffered to indicated pH and supplemented with 7.5 mM GlcNAc and 0.05% casamino acids
[b] *p*-values and fold changes compare to wild type at pH 8.5; $p > 0.05$ for Δ*aphB* vs. wild type at pH 5.5
[c] Cultures grown in SWT
[d] Cultures grown in LB

**Table S3. Bacterial strains and plasmids used in this work**

| Strain | Description | Source |
|---|---|---|
| ES114 | *Vibrio fischeri*, wild-type | (1) |
| BDB127 | ES114 with empty pVSV105 | This work |
| BDB128 | ES114 with pVSV105::*hbtRC* | This work |
| BDB171 | ES114 with pVSV105::*litR* | This work |
| BDB156 | ES114 with pVSV102 | (2) |
| BDB011 | ES114 with pVSV208 | (2) |
| CBNR46 | ES114 *att*Tn*7*:: $P_{lac}$-*gfp-erm* | (3) |
| CBNR47 | ES114 *att*Tn*7*:: $P_{lac}$-*rfp-erm* | (3) |
| BDB003 | ES114 Δ*hbtRC* (VF_A0473–A0474) | (4) |
| BDB134 | ES114 Δ*hbtRC* (VF_A0473–A0474), newly derived | This work |
| BDB023 | Δ*hbtRC* (BDB003) with pVSV105::*lacI-hbtRC* | This work |
| BDB039 | Δ*hbtRC* (BDB003) with empty pVSV105 | This work |
| BDB139 | Δ*hbtRC* (BDB134) with empty pVSV105 | This work |
| BDB140 | Δ*hbtRC* (BDB134) with pVSV105::*hbtRC* | This work |
| BDB154 | Δ*hbtRC* (BDB134) with pVSV105::*tcpPH* | This work |
| BDB157 | Δ*hbtRC* (BDB134) with pVSV102 | This work |

| | | |
|---|---|---|
| BDB012 | Δ*hbtRC* (BDB134) with pVSV208 | This work |
| BDB078 | Δ*hbtRC* (BDB003) *att*Tn*7*:: $P_{lac}$-*gfp-erm* | This work |
| BDB079 | Δ*hbtRC* (BDB003) *att*Tn*7*:: $P_{lac}$-*rfp-erm* | This work |
| BDB172 | Δ*hbtRC* Δ*toxRS* (VF_0790–91) | This work |
| BDB173 | Δ*hbtRC* Δ*toxRS* with empty pVSV105 | This work |
| BDB174 | Δ*hbtRC* Δ*toxRS* with pVSV105::*hbtRC* | This work |
| BDB182 | ES114 Δ*litR* (VF_2177) | This work |
| BDB183 | Δ*litR* with empty pVSV105 | This work |
| BDB184 | Δ*litR* with pVSV105::*hbtRC* | This work |
| BDB185 | Δ*litR* with pVSV105::*litR* | This work |
| BDB129 | ES114 ΔVF0157–0180 | This work |
| BDB186 | ΔVF0157–0180 with empty pVSV105 | This work |
| BDB187 | ΔVF0157–0180 with pVSV105::*hbtRC* | This work |
| BDB188 | ΔVF0157–0180 with pVSV105::*litR* | This work |
| BDB135 | ES114 ΔVF_2042 | This work |
| BDB141 | ES114 ΔVF_1133 | This work |
| BDB142 | ES114 ΔVF_A0389 | This work |
| BDB144 | ES114 ΔVF_A0246 | This work |

| | | |
|---|---|---|
| BDB163 | ES114 ΔVF_1133 ΔVF_2042 ΔVF_A0246 ΔVF_A0389 | This work |
| BDB130 | ES114 Δ*aphB* (VF_1690) | This work |
| JB24 | Δ*crp* (VF_2280) | (5) |
| CA1 | Δ*rpoQ* (VF_A1015) | (6) |
| BDB194 | Δ*rpoQ* with empty pVSV105 | This work |
| BDB195 | Δ*rpoQ* with pVSV105::*hbtRC* | This work |
| BDB176 | Δ*rpoQ* with pVSV105::*litR* | This work |
| 0395-N1 | *V. cholerae* Ogawa 395 Δ*ctxA* | (7) |
| BDB190 | 0395-N1 Δ*tcpPH* (ΔVC0395_A0351-A0352) | This work |
| BDB191 | Δ*tcpPH* with empty pVSV105 | This work |
| BDB192 | Δ*tcpPH* with pVSV105::*hbtRC* | This work |
| BDB193 | Δ*tcpPH* with pVSV105::*tcpPH* | This work |
| DH5α*pir* | *E. coli* cloning strain; *recA1 endA1 hsdR17 supE44 thi-1 Δ(lacZYA-argF)U169 [Φ80dlacZΔM15] gyrA96 relA1 λpir+* | (8) |
| WM3064 | *E. coli* conjugation strain; *thrB1004 pro thi rpsL hsdS lacZΔM15 RP4-1360 Δ(araBAD)567 ΔdapA1341::[erm pir(wt)]* | (9) |

| Plasmid | Description | Source |
|---|---|---|
| pAKD601 | IPTG-inducible expression vector | (2) |
| pVSV105 | Cloning vector; $Cm^r$ | (2) |
| pVSV105::*lacI-hbtRC* | *lacI^q^-A1*/*O4*/*O3*p from pAKD601, 26 bp upstream of *lacI^q^*, 7 bp downstream of *A1*/*O4*/*O3*p; *hbtRC*, 18 bp upstream, 48 bp downstream; $Cm^r$ | This work |
| pVSV105::*hbtRC* | *hbtRC*, 18 bp upstream, 48 bp downstream; $Cm^r$ | This work |
| pVSV105::*litR* | *litR*, 21 bp upstream, 87 bp downstream; $Cm^r$ | This work |
| pVSV105::*tcpPH* | *tcpPH*, 20 bp upstream, 2 bp downstream; $Cm^r$ | This work |
| pVSV102 | *gfp*; $Km^r$ | (2) |
| pVSV208 | *rfp*; $Cm^r$ | (2) |
| pEVS107 | mini-Tn7 vector; $Km^r$, $Em^r$ | (10) |
| pCBNR6 | pEVS107::$P_{lac}$-*gfp*; $Km^r$, $Em^r$ | (3) |
| pCBNR7 | pEVS107::$P_{lac}$-*rfp*; $Km^r$, $Em^r$ | (3) |
| pUX-BF13 | *tnsABCDE* transposase vector | (11) |
| pEVS104 | Conjugation helper; $Km^r$ | (12) |

| | | |
|---|---|---|
| pSMV3 | Deletion vector, Km$^r$, *sacB* | (13) |
| pSMV3Δ*hbtRC* | pSMV3 with VF_A0473–A0474 flanking sequences | This work |
| pSMV3Δ*litR* | pSMV3 with VF_2177 flanking sequences | This work |
| pSMV3ΔVF_0157-80 | pSMV3 with VF_0157-80 flanking sequences | This work |
| pSMV3ΔVF_1133 | pSMV3 with VF_1133 flanking sequences | This work |
| pSMV3ΔVF_2042 | pSMV3 with VF_2042 flanking sequences | This work |
| pSMV3ΔVF_A0246 | pSMV3 with VF_A0246 flanking sequences | This work |
| pSMV3ΔVF_A0389 | pSMV3 with VF_A0389 flanking sequences | This work |
| pSMV3 ΔVC0395_ A0351-A0352 | pSMV3 with VC0395_A0351-A0352 flanking sequences | This work |

**Table S4. Primers and probes used in this work**

| Primer | Sequence | Restriction Site |
|---|---|---|
| <u>*lacI* and *hbtRC* insertion into pVSV105</u> | | |
| lacIF2 | CTAGGTCGACGCTAACTTACATTAATTGCGTTG | *Sal*I |
| lacIR | CTAGGAATTCCTGTGTGAAATTGTTATCCGC | *Eco*RI |
| hbtRF | CTAGGAATTCCTGTTAAGTCAGGATGATATGG | *Eco*RI |
| hbtCR | CATGGGTACCGTACCCTAAACACACTCTATAAATAAC | *Kpn*I |
| <u>*hbtRC* insertion into pVSV105</u> | | |
| hbtRF2 | CTAGTCTAGACTGTTAAGTCAGGATGATATGG | *Xba*I |
| hbtCR | CATGGGTACCGTACCCTAAACACACTCTATAAATAAC | *Kpn*I |
| <u>*litR* insertion into pVSV105</u> | | |
| litRF | GTACGTCGACGTTGGCAAGGATATAAATATAATGG | *Sal*I |
| litRR | GTACGGTACCTTCTGATTAACAACGCATTTG | *Kpn*I |
| <u>*tcpPH* insertion into pVSV105</u> | | |
| VCtcpPF | CATGTCTAGAGATTAAGAAAATGTAAAGTAATGG | *Xba*I |
| VCtcpHR | CATGGGTACCCCCTAAAAATCGCTTTGACAG | *Kpn*I |
| <u>Gene deletion</u> | | |
| hbtRUSF | GATCGGATCCGAGAGCATTAAGTAGTGTTAATC | *Bam*HI |
| hbtRUSR | GATCGGTACCCCATATCATCCTGACTTAACAG | *KpnI* |
| hbtCDSF | GATCGGTACCCAGCAGGAAGTGGTAGTCTC | *Kpn*I |
| hbtCDSR | GATCGAGCTCCAGGACTATTCGGGTTAAG | *Sac*I |
| litRUSF | GATCGGATCCGCATGATTAAGCGTTAAGATTAG | *Bam*HI |

| | | |
|---|---|---|
| litRUSR | GATCGAATTCCCATTATATTTATATCCTTGCCAAC | *Eco*RI |
| litRDSF | GATCGAATTCCAAATTGTTTCTTAAATATGTTGTG | *Eco*RI |
| litRDSR | GATCGAGCTCGGTGGTATCATCGGTCTTG | *Sac*I |
| VF1133USF | GTACGGATCCGGTCACTTTGCACTCTCAC | *Bam*HI |
| VF1133USR | GTACGGTACCCTGTTCATGTTGTATCTCTCTTTC | *Kpn*I |
| VF1133DSF | GTACGGTACCGTAATTATCTTAGATAAGCAATGTC | *Kpn*I |
| VF1133DSR | GTACGAGCTCGCGTTTGGCATTGAGAATTG | *Sac*I |
| VF2042USF | GTACGAATTCGAAACCGTTTCAATCAAAACAG | *Eco*RI |
| VF2042USR | GTACGGTACCCATAGCTGCATACCTTAGTTATAAC | *Kpn*I |
| VF2042DSF | GTACGGTACCCTGTAATTAAGTTGATTTTTCACTTATG | *Kpn*I |
| VF2042DSR | GTACCTCGAGCCAGCTTTAATTGAGCCTTC | *Xho*I |
| VFA0246USF | GTACGGATCCCGCCTAATGCCTTCAATCG | *Bam*HI |
| VFA0246USR | GTACGGTACCCATAGATAACACCGCTGTAAG | *Kpn*I |
| VFA0246DSF | GTACGGTACCGTATAAAGGGTAATTCAATAGGCTG | *Kpn*I |
| VFA0246DSR | GTACGAGCTCCCAAGCCATCTCAATTTTACG | *Sac*I |
| VFA0389USF | GTACGGATCCGCCAGCCTTATCAGTTTTATTG | *Bam*HI |
| VFA0389USR | GTACGAATTCCTTCATAGAAACATCCTAAATCCG | *Eco*RI |
| VFA0389DSF | GTACGAATTCGTGTAACTGACGTATAACTATC | *Eco*RI |
| VFA0389DSR | GTACGAGCTCGCGTTACGACTATTGGTTTAC | *Sac*I |
| VF0157USF | GTACGGATCCGTCTAAAAATAAGAAAGCAGTGC | *Bam*HI |
| VF0157USR | GTACGGTACCGAAAAGGGCTTTGAGGGG | *Kpn*I |
| VF0180DSF | GTACGGTACCGATAAGCAAATAATGGCGTTTTG | *Kpn*I |
| VF0180DSR | GTACGAGCTCGAGTAACTTCGCTTAATGCC | *Sac*I |

| | | |
|---|---|---|
| VF1690USF | GTACGGATCCCTGATCAATAAACAAGAGATAACG | *Bam*HI |
| VF1690USR | GTACGGTACCCTAATTTCATTATATTAAGATACTCTCC | *Kpn*I |
| VF1690DSF | GTACGGTACCGTAACCACATACCTGATTAATTTAC | *Kpn*I |
| VF1690DSR | GTACgagctcGTTTAGGTAAGCGTGAAATTG | *Sac*I |
| toxRUSF | GATCGGATCCGAACTCTTGGTATTCAGCATC | *Bam*HI |
| toxRUSR | GATCGGTACCCATCGGTTATTAACTCAGTAGTG | *Kpn*I |
| toxSDSF | GATCGGTACCCTATTTTCGCATTAACAATGATTTC | *Kpn*I |
| toxSDSR | GATCGAGCTCGAAATTTGGAAAAGATGACAGG | *Sac*I |

Expression analysis

| | |
|---|---|
| VF1133expF | CAACAAAGCGTCACTCAAAC |
| VF1133expR | CCTCTAAGCTACGTGCAATC |
| VF2042expF | CAGGTGAACAAGGTCGAG |
| VF2042expR | CCTTCTTGGCAATCACGAC |
| VFA0246expF | GGTGCAGAGTGAAGCTAATG |
| VFA0246expR | TGAACCTTCAACGGTTAGTG |
| VFA0389expF | GAACGTAGCATCAGGAGAAG |
| VFA0389expR | GTGACTTAACGCTTTGGTTTC |
| flaAexpF | GTCTGTAGGTGATGCACAAG |
| flaAexpR | CGGCTGTTAGAAGCAGATAC |
| flaCexpF | GGTGGTTCTCGTCTTCTTAATG |
| flaCexpR | TCAGCATTTACAGCCCAATC |
| flaFexpF | GGAAAGTAGGTGCAGAGAATAC |
| flaFexpR | CGTCTCACCTGACATGAATAC |

| | |
|---|---|
| motA1expF | CCTGCAATGGGTATGATTGG |
| motA1expR | TGAGCAATAGGGAATGCAAC |
| flgMexpF | CCAGAAACTCAAGCACCTG |
| flgMexpR | CGTTCAGGATCAACTGTGTATG |
| flrAexpF | TCGTGCAGATGTTCGTATTG |
| flrAexpR | CTGCCTTCAGCTTCCATTC |
| flrBexpF | ACCGCTCTGCAATCAAATC |
| flrBexpR | TGACATCCCTTCCCTATCAC |
| VF0157expF | CCAAATGTACGCTGGCTATC |
| VF0157expR | CCATAATCACCGCCTTCTTG |
| VF0164expF | GATTCTGCTGGTGTTGCG |
| VF0164expR | GGCAAACGTCACTGGTTAG |
| VF0165expF | GTACCAGTATGCACTTTGCC |
| VF0165expR | CGGCATCCACAACCAATAC |
| hbtRexpF | GCTGTTCCTGCAACAAGTAG |
| hbtRexpR | GAGACTACCACTTCCTGCTG |
| litRexpF | GGAGTACCTCTACTCGTG |
| litRexpR | CAGGGGTATGATCGTTAC |
| toxTexpF | AAAGAACTAGAGTCTCGAGGAG |
| toxTexpR | ACTTGGTGCTACATTCATGG |
| VCpolAqF | AAGAGCTGGCTTTGGATTAC |
| VCpolAqR | CCGTCACTGCTTGATGATAG |

*In situ* HCR probes targeting *hbtR*

ATGGTTTTTAAACTCAATGAAATCTATTGGGATCCAGCAACAAAAAAATTGT

GATACGCTAGAAGATGCAGTTAATGGTAATAACGAATATGGTTCAACAACGC

TTAGTTTCTGCAATTCTGACTTTGTTAATAAAAGAACATCCTTCTATTTGTA

AATGAACATATTAAAGATGCTCTTTGGGGGACTCAGTGGATATCAAATGAAA

ATTCCTCAGTTAATTAAAAGAACAAGAGTATCAATAAAAGATACGAATAGAC

GTTATCGAAAATGTAAAAGGCAATGGTTATAAAATAAACAATGTAGAAGAGA

AGAATAAATTCAAATAGCATACAATGCACAGAGAAAAAAACGTCACATTCTA

TCAGCATTAATCTTTTGTATAGAAAAACATGTGTTTTACGATCTAATTCCTT

GAAGAGATAAAAAAAGTCAAAGATTTTGATTTAATTAAAATCTCTGAAGATA

TTTTTAATTAAGACTAAAGACCAAGAGTGTGAATTGAACCTTAAAAATAAAA

*In situ* HCR probes targeting *litR*

ATGGATACAATCCAAAAAAGGCCTAGAACAAGGCTATCTCCAGAAAAGCGCA

GAACAATTACTTGATATCGCAATTGAAGTGTTTTCACAACGTGGCATTGGTC

GGTGGTCATGCTGATATTGCAGAAATCGCTCAGGTCTCTGTCGCTACCGTGT

AATTACTTCCCAACAAGAGAAGATCTTGTTGATGATGTATTAAATAAAGTCG

AACGAATTTCACCAATTCATCAATAACTCAATTTCTCTTGATCTTGATGTTC

TCTAACCTTAATACTCTTCTTCTGAATATCATTGACAGTGTTCAAACTGGTA

AAGTGGATCAAAGTATGGTTTGAATGGAGTACCTCTACTCGTGATGAAGTAT

CCTCTTTTCTTAAGTACTCACTCAAATACAAATCAAGTTATCAAAACTATGT

GAAGAAGGTATTGAACGTAACGAAGTATGTAACGATCATACCCCTGAAAACC

ACAAAAATGCTTCATGGTATTTGTTACTCTGTTTTCATTCAGGCTAACCGTA

AGTTCATCTGAAGAAATGGAAGAAACGGCAAATTGTTTCTTAAATATGTTGT

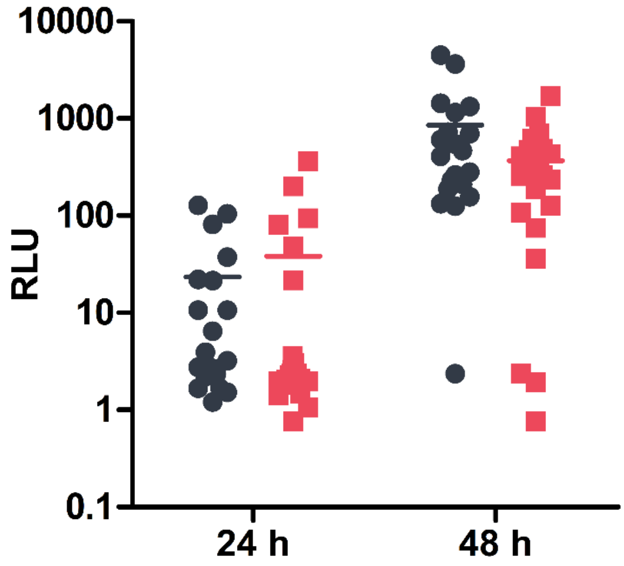


**FIG S1** Luminescence of wild-type *V. fischeri* and Δ*hbtRC* strains during symbiosis. Luminescence was measured in juvenile squid 24 h or 48 h after colonization with either wild-type *V. fischeri* (●) or Δ*hbtRC* (■). Each point represents one animal. Mean values indicated; no significant differences were noted between strains at either time-point. RLU, relative light units.

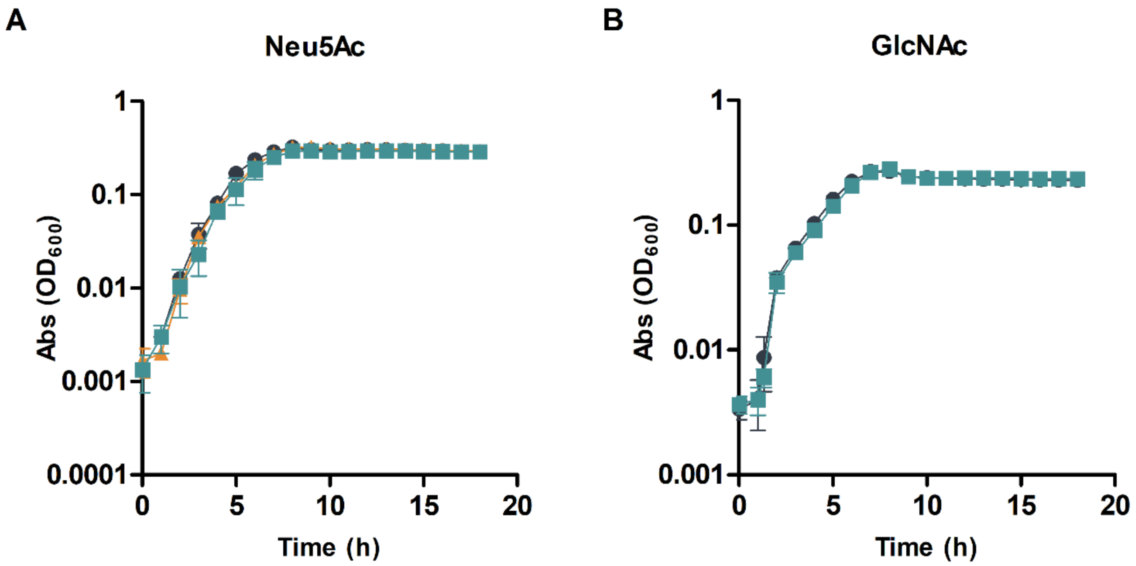


**FIG S2** Growth of wild-type *V. fischeri* and chemotaxis mutants on *N*-acetylated sugars. (A) The rates of growth in MSM supplemented with 0.05% casamino acids and 6.5 mM Neu5Ac were measured for wild-type *V. fischeri* (●), ΔΔΔΔ (■), and ΔVF_1133 ΔVF_A0246 (▲). (B) The rates of growth in MSM supplemented with 0.05% casamino acids and 6.5 mM GlcNAc were measured for wild-type *V. fischeri* (●) and ΔΔΔΔ (■). Results represent means of three biological replicates ± one standard deviation. Abs, absorbance.

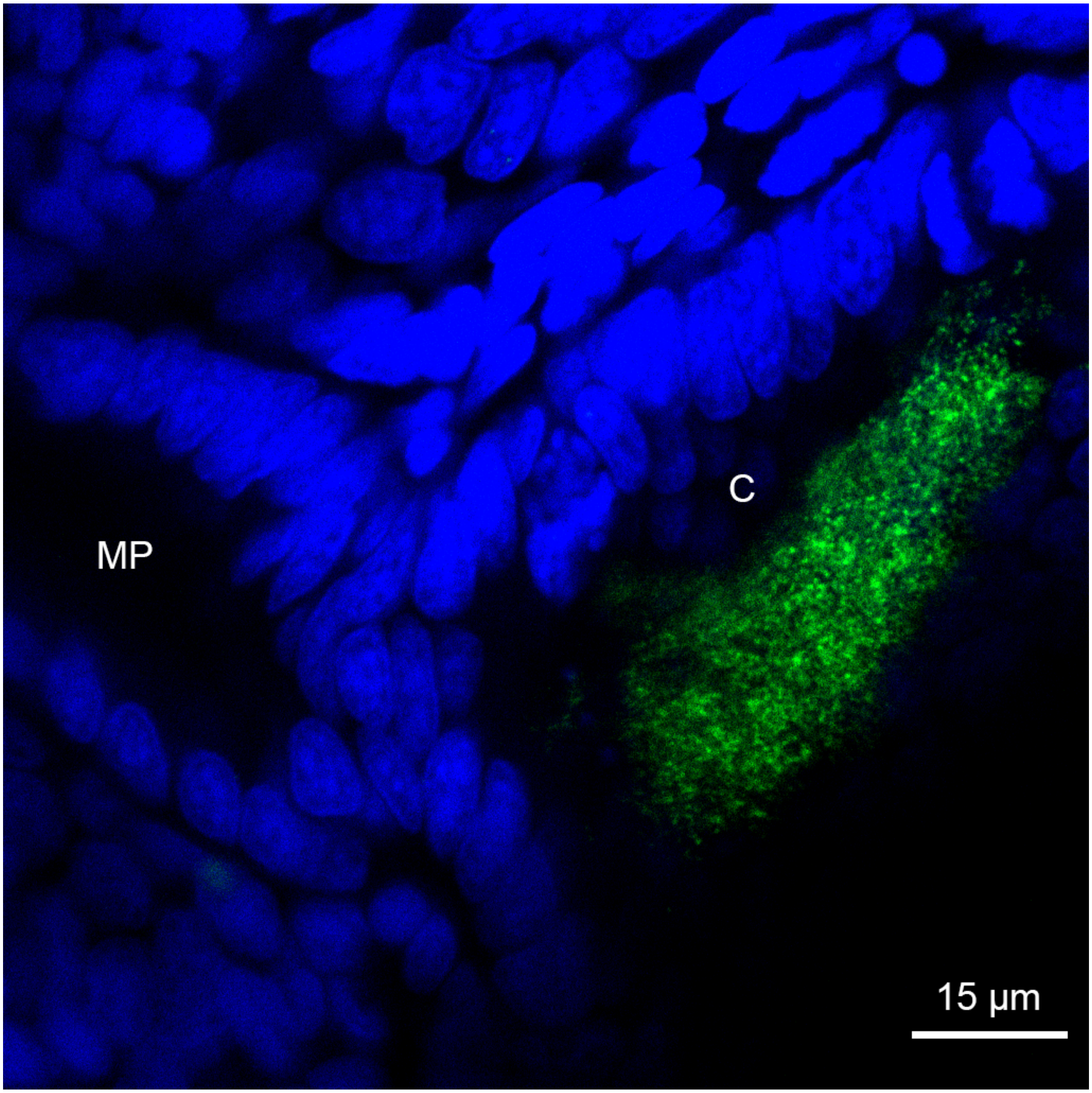


**FIG S3** Expression of *litR* in *V. fischeri* when host-associated within the *E. scolopes* light organ. Transcripts of *litR* (green) were detected using *in situ* HCR in *V. fischeri* cells colonizing *E. scolopes* crypts (C) after traversing the tissues of the migration path (MP). Blue, *E. scolopes* nuclei (DAPI); green, *litR* transcripts (Alexa 546). Bar 15 µm.
